# A data-driven framework for the selection and validation of digital health metrics: use-case in neurological sensorimotor impairments

**DOI:** 10.1101/544601

**Authors:** Christoph M. Kanzler, Mike D. Rinderknecht, Anne Schwarz, Ilse Lamers, Cynthia Gagnon, Jeremia Held, Peter Feys, Andreas R. Luft, Roger Gassert, Olivier Lambercy

## Abstract

**Background:** Digital health metrics have the potential to advance the monitoring and understanding of impaired body functions, for example in persons with neurological disorders. However, their integration into clinical research and practice is challenged by insufficient validation of the vast amount of existing and often abstract metrics. Here, we propose a data-driven framework to select and validate a clinically-relevant core set of digital health metrics extracted from a technology-aided assessment. As a use-case, this framework is applied to metrics extracted from the Virtual Peg Insertion Test (VPIT), a sensor-based assessment of upper limb sensorimotor impairments.

**Methods:** The framework builds on a use-case specific pathophysiological motivation of digital health metrics to represent clinically-relevant impairments, models the influence of confounds from participant demographics, and evaluates the most important clinimetric properties (discriminant validity, structural validity, test-retest reliability, measurement error, learning effects). This approach was applied to 77 kinematic and kinetic metrics extracted from the VPIT, using data from 120 neurologically intact controls and 89 subjects with neurological disorders (post-stroke, multiple sclerosis, or hereditary ataxia). An exploratory factor analysis to discuss the initially proposed pathophysiological hypotheses was performed and the sensitivity of the metrics to clinically-defined disability levels was investigated.

**Results:** Applied to the VPIT, the framework selected 10 (13.0%) clinically-relevant core metrics. These assess the severity of multiple sensorimotor impairments in a valid, reliable, and informative manner for all three disorders while being least susceptible to measurement error and learning effects. The digital health metrics of the VPIT provided additional clinical value by detecting impairments in neurological subjects that did not show any deficits according to conventional scales, and by covering several sensorimotor impairments of the arm and hand with a single assessment.

**Conclusions:** The proposed framework could help to address the insufficient evaluation, standardization, and interpretability of digital health metrics. In the presented use-case, it allowed to establish validated core metrics for the VPIT, paving the way for its integration into clinical neurorehabilitation trials.

## 1 Introduction

Assessments of impaired body functions, as observed in many diseases and disorders, are a fundamental part of the modern healthcare system [1]. Specifically, these assessments are essential to provide documentation for insurances, to individualize therapeutic interventions, and to shed light on the often unknown mechanisms underlying the impairments and their temporal evolution. An exemplary application scenario of assessments are neurological disorders, including stroke, multiple sclerosis (MS), and hereditary ataxic conditions, where impairments in the sensorimotor system are commonly present, for example when coordinating arm and hand during goal-directed activities [2–5]. In research studies, such deficits are often assessed by healthcare practitioners, who subjectively evaluate persons with impairments during multiple standardized tasks (referred to as *conventional scales*) [6–8]. While most of these scales are validated and their interpretation fairly well understood and documented, they often have a limited ability to detect fine impairments because of limited knowledge about behavioral variability, low resolution, and ceiling effects, leading to bias when attempting to model and better understand longitudinal changes in impairment severity [9–12].

Digital health metrics, herein defined as discrete one-dimensional metrics that are extracted from health-related sensor data, promise to overcome these shortcomings by proposing objective and traceable descriptions of human behaviour without ceiling effects and with high resolution [13]. This offers the potential to more sensitively characterize impairments and significantly reduce sample sizes required in resource-demanding clinical trials [14]. In the context of assessing sensorimotor impairments, a variety of digital health metics relying on kinematic or kinetic data have been successfully applied to characterize abnormal movement patterns [13, 15, 16]. However, the integration of digital health metrics into clinical routine and research is still inhibited by an insufficient evaluation of the vast amount of existing measures and the need for core sets of validated and clinically-relevant measures for the targeted impairments [13, 17–20]. Indeed, recent reviews reported the use of over 150 sensor-based metrics for quantifying upper limb sensorimotor impairments and highlighted a clear lack of evidence regarding their pathophysiological motivation and clinimetric properties [13, 21]. Especially the ability of a metric to detect impairments (discriminant validity) as well as the dependency to other metrics and the underlying information content (structural validity) are often not evaluated. Similarly, test-retest reliability, measurement error arising from intra-subject variability, and learning effects are only rarely considered, but their evaluation is fundamental to reliably and sensitively quantify impairments in an insightful manner [22]. Further, the influence of participant demographics, such as age, sex, and handedness, on the metrics is often not accurately modeled, but needs to be taken into account to remove possible confounds and provide an unbiased assessment. Most importantly, the high variability of clinimetric properties across behavioral tasks and sensor-based metrics motivates the need for a methodology to select metrics for a specific assessment task, starting from a large set of potential metrics that should be narrowed down to a clinically-relevant core set [13, 17, 23]. Unfortunately, existing approaches to select core sets often do not consider the pathophysiological interpretation of metrics or are rarely tailored to the specific requirements of digital health metrics (e.g,. sufficient clinimetric properties) [17, 24–28].

Hence, the objective of this work was to propose and apply a data-driven framework to select and validate digital health metrics, aimed at providing evidence that facilitates their clinical integration. The approach relies on i) a use-case specific pathophysiological motivation for sensor-based metrics to represent clinically-relevant impairments, considers ii) the modeling of confounds arising through participant demographics, and implements iii) data processing steps to quantitatively evaluate metrics based on the most important clinimetric properties (discriminant validity, structural validity, test-retest reliability, measurement error, and learning effects). Herein, we present this framework in the context of a use-case with the Virtual Peg Insertion Test (VPIT), an instrumented assessment of upper limb sensorimotor impairments consisting of a goal-directed manipulation task in a virtual environment [29–34]. For this purpose, 77 kinematic and kinetic metrics were extracted from VPIT data from a cohort of neurologically intact and affected subjects (stroke, MS, and hereditary ataxia). We hypothesized that the presented methodology would be able to reduce a large set of metrics to a core set with optimal clinimetric properties that allows assessing the severity of the targeted impairments in a robust and insightful manner.

Targeting this objective is important, as the proposed data-driven framework can easily be applied to metrics gathered with other digital health technologies. This will help addressing the lacking evaluation, standardization, and interpretability of digital health metrics, a necessary step to address their still limited clinical relevance [19, 20, 35]. Further, the presented use-case establishes a validated core set of metrics for the VPIT, paving the way for its integration into clinical trials in neurorehabilitation.

## 2 Methods

To objectively reduce a large set of digital health metrics to a clinically-relevant subset, we implemented a three-step process (Figure 1) considering the most important statistical requirements to sensitively and robustly monitor impairments in a longitudinal manner. These requirements were inspired from the COSMIN guidelines for judging the quality of metrics based on systematic reviews and related work on digital health metrics [13, 22, 36–38]. Further, two additional validation steps were implemented to improve the understanding of the selected core metrics (Figure 1). While this selection and validation framework is independent of a specific assessment platform (i.e., the initial set of metrics to be evaluated), the manuscript defines the framework in the context of the VPIT with the goal to provide specific instructions including a hands-on example, starting from the initial motivation of metrics to the selection of a validated core set.

**Figure 1:**
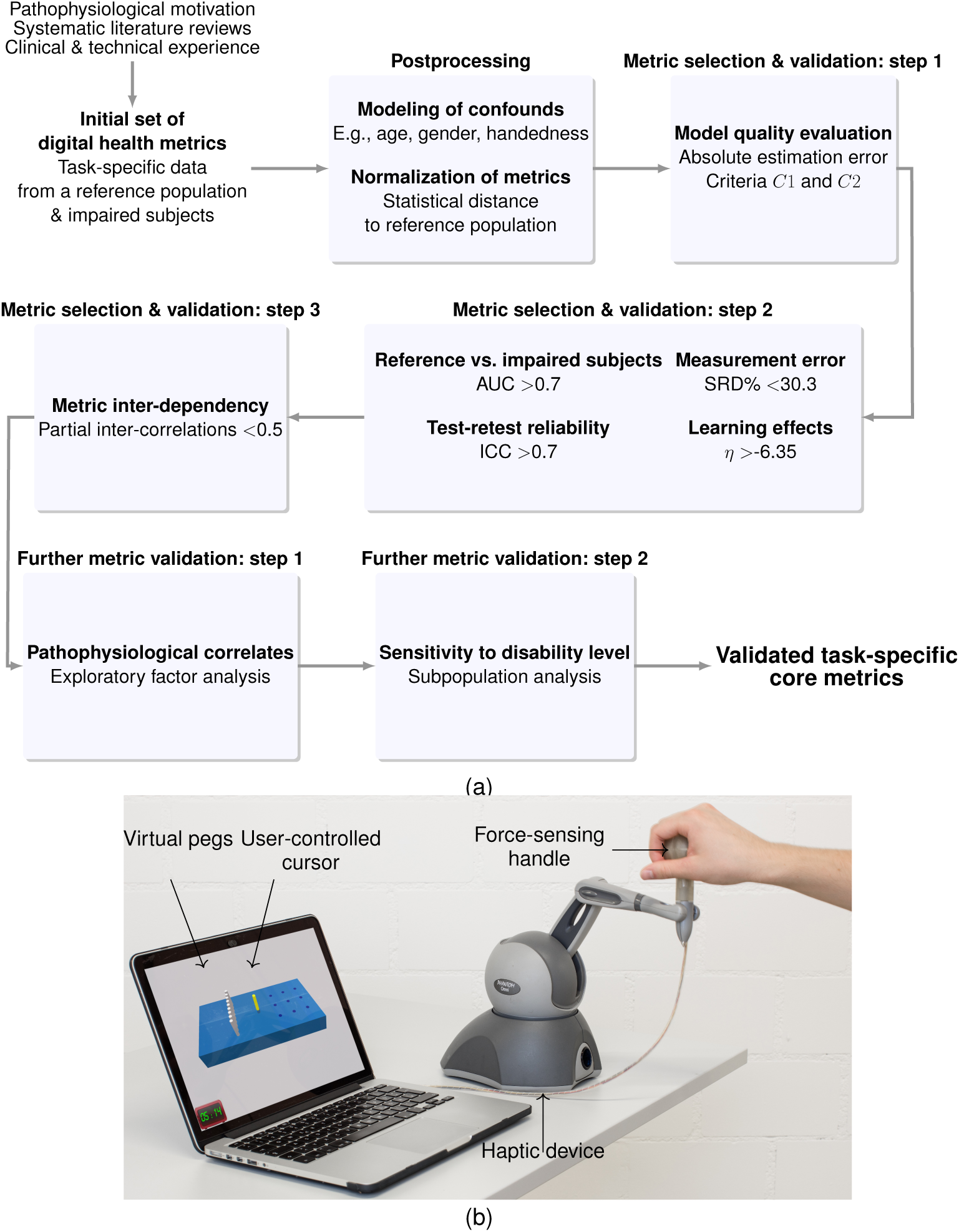
Overview of the data-driven framework and the Virtual Peg Insertion Test (VPIT). (a) The frameworks allows to select a core set of validated digital health metrics. Criteria *C*1 and *C*2 defining model quality; ROC: receiver operating characteristics; AUC: area under curve; ICC: intra-class correlation; SRD%: smallest real difference; *η* strength of learning effects; (b) as a use-case, the framework was applied to data recorded with the VPIT, a sensor-based upper limb sensorimotor assessment requiring the coordination of arm and hand movements as well as grip forces. The test combines a commercial haptic device, a handle instrumented with force sensors, and a virtual pegboard.

### 2.1 Virtual Peg Insertion Test

The VPIT is a digital health assessment combining a commercial haptic end-effector (PHANTOM Omni/Touch, 3D Systems, CA, USA), a custom-made handle with piezoresistive force sensors (CentoNewton40, EPFL, Switzerland), and a virtual reality (VR) environment, implemented in C++ and OpenGL on a Microsoft (Redmond, WA, USA) Windows laptop (Figure 1). The assessment features a goal-directed pick-and-place task that requires arm and hand movements while actively lifting the arm against gravity, thereby combining elements of the Nine Hole Peg Test (NHPT) and the Box and Block Test [39, 40]. The VR environment displays a rectangular board with nine cylindrical pegs and nine corresponding holes arranged as a 3×3 matrix with the same dimensions as the NHPT (31.1 26.0 4.3 cm) [39]. The objective is to transport the virtual pegs into the holes by controlling a cursor through the 6D-movements (3D-position and 3D-angular orientation) of the haptic device, which provides up to 3.3 N of haptic feedback to render the virtual pegboard. A peg can be picked up by aligning the position of a cursor with the peg (alignment tolerance: 3.0 mm) and applying a grasping force above a 2 N threshold. The peg needs to be transported towards a hole while maintaining a grasping force of at least 2 N, and can be inserted in the hole by releasing the force below the threshold, once properly aligned with a hole. The holes in the board of the VR environment are rendered through reduced haptic impedance compared to other parts of the board. The pegs cannot be picked up anymore upon insertion in a hole and are perceived as transparent throughout the test (i.e., no collisions between pegs are possible). The default color of the cursor is yellow and changes after spatially aligning cursor and peg (orange), during the lifting of a peg (green), or after applying a grasping force above the threshold while not being spatially aligned with the peg (red). During the execution of the task, 6D-endpoint position, grasping forces, and interaction forces with the VR environment are recorded at 1 kHz.

### 2.2 Participants & procedures

The analysis presented in this work builds on data from different studies that included assessments with the VPIT [30, 41–43]. In addition, age-matched reference data was based on 120 neurologically intact subjects. Their handedness was evaluated using the Edinburgh Handedness Inventory and potential stereo vision deficits that might influence the perception of a virtual environment were screened using the Lang stereo test [44]. Sixty of these subjects were further tested a second time one to three days apart to evaluate test-retest reliability. Additionally, 53 post-stroke subjects, 28 MS subjects, and 8 subjects with autosomal recessive spastic ataxia of Charlevoix-Saguenay (ARSACS) were tested. Each subject was tested with the VPIT on both body sides if possible. The administered conventional assessments were dependent on the disease and the specific study. Commonly applied assessments were the Fugl-Meyer upper extremity (FMA-UE) [9], the Nine Hole Peg Test (NHPT) [39], and the Action Research Arm Test (ARAT) [45]. Detailed exclusion criteria and ethical approval references are listed in the supplementary material (SM). All subjects gave informed written consent.

To perform the VPIT, participants were seated in a chair with backrest and without armrests in front of a personal computer with the haptic device being placed on the side of the tested limb. The initial position of the subjects (i.e., hand resting on the handle) was defined by a shoulder abduction angle of 45*^◦^*, a shoulder flexion angle of 10*^◦^*, and an elbow flexion angle of 90*^◦^*. Subjects were familiarized with the task and subsequently performed five repetitions (i.e., inserting all nine pegs five times) per body side. Participants were instructed to perform the task as fast and accurately as possible.

### 2.3 Data preprocessing

Data preprocessing steps are required to optimize the quality of the sensor data and dissect the complex recorded movement patterns into distinct movement phases that can be related to specific sensorimotor impairments. First, temporal gaps larger than 50 samples in the recorded position, force, and haptic time-series were linearly interpolated. Such gaps can stem from a delayed communication between the soft- and hardware components during the data recordings. Subsequently, a 1D distance trajectory *d*(*t*) was estimated from the 3D cartesian position trajectories *p_x_*, *p_y_*, and *p_z_* by summing up their absolute first time-derivatives until timepoint *t*:

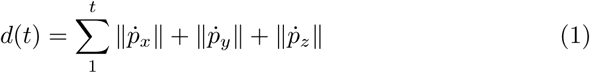

Afterwards, velocity (first time-derivative) and jerk (third time-derivative) signals were derived from *d*(*t*). Also, single grasping force and grip force rate (first time-derivative) trajectories were generated by averaging across the signals of the three piezoresistive sensors. All time-series were low-pass filtered initially and after each derivation using a zero-phase Butterworth filter (4*^nd^* order, cut-off frequency 8 Hz). Data from an entire peg were removed if it was dropped and not inserted into a hole before another peg was picked up.

To isolate rapid ballistic movements, the trajectories of each peg were segmented into the *transport* (i.e., ballistic movement while transporting the peg to a hole) and *return* (i.e., ballistic movement while returning the cursor to the next peg) phases (Figure SM1). The *transport* phase started at the last occasion the velocity exceeded a threshold *θ_vel,tp_* after the peg was picked up and before maximum velocity *v_max,tp_* was reached. The threshold *θ_vel,tp_* was set to 10% of *v_max,tp_* that occurred before the insertion of the peg into the next hole. The end of the *transport* was defined as the first time the velocity dropped below *θ_vel,tp_* after *v_max,tp_*. To ensure a robust segmentation, the *transport* phase of a peg was discarded in case the peg was taken at *v_max,tp_*, the velocity never dropped below *θ_vel,tp_* after *v_max,tp_* before releasing the peg, or the length of the phase was below 0.1 s. The same criteria were applied to segment the *return* phase, which was defined as the main ballistic movement component between releasing a peg and picking up the next peg, given the maximal velocity *v_max,rt_* during return and *θ_vel,rt_*. For segmenting the *transport* and *return* phases, only the horizontal component of *d*(*t*) was used [46].

To isolate the overshoot when reaching for a target as well as the precise position adjustments related to virtual object manipulations, the trajectories were additionally segmented into the *peg approach* and *hole approach* phases. The former was defined from the end of the *return* until the next peg was picked up. The latter was defined from the end of the *transport* until the current peg was inserted into a hole.

Further, grasping forces were additionally segmented into the *force buildup* (i.e., behaviour during the most rapid production of force) and *force release* phases (i.e., behaviour during the most rapid release of force), by first identifying the position of the maximum and minimum value in grip force rate between approaching and inserting each peg (Figure SM1). Subsequently, the start and end of the *force buildup* phase was defined as the last and first time the grip force rate was below 10% of its maximum before and after the maximum, respectively. Similarly, the start and end of the *force release* phase was determined based on the last and first time the grip force rate was above 10% of its minimum value before and after the minimum, respectively.

### 2.4 Pathophysiological motivation of digital health metrics

To facilitate the pathophysiological interpretation of sensor-based metrics for each use-case, it is of importance to describe the mechanisms underlying a specific disease, their effect on the assessed behavioral construct, and how metrics are expected to capture these abnormalities. Within the use-case of the VPIT, this pathophysiological motivation is implemented using the computation, anatomy, and physiology model as well as the clinical syndromes ataxia and paresis that are commonly present in neurological disorders [47, 48]. Leveraging these concepts allows to especially connect how inappropriately scaled motor commands and an inability to voluntarily activate spinal motor neurons affect upper limb movement behaviour. As the VPIT strives to capture multiple heterogeneous and clinically-relevant sensorimotor deficits, a variety of different movement characteristics were defined to describe commonly observed upper limb sensorimotor impairments in neurological disorders. Subsequently, an initial set of 77 metrics (Table 1 and 2) for the VPIT were proposed with the aim to describe these movement characteristics and the associated sensorimotor impairments. These metrics were preselected based on the available sensor data (i.e., end-effector kinematic, kinetics, and haptic interactions), recent systematic literature reviews as well as evidence-based recommendations [13, 21, 49], and the technical and clinical experience of the authors.

**Table 1:**
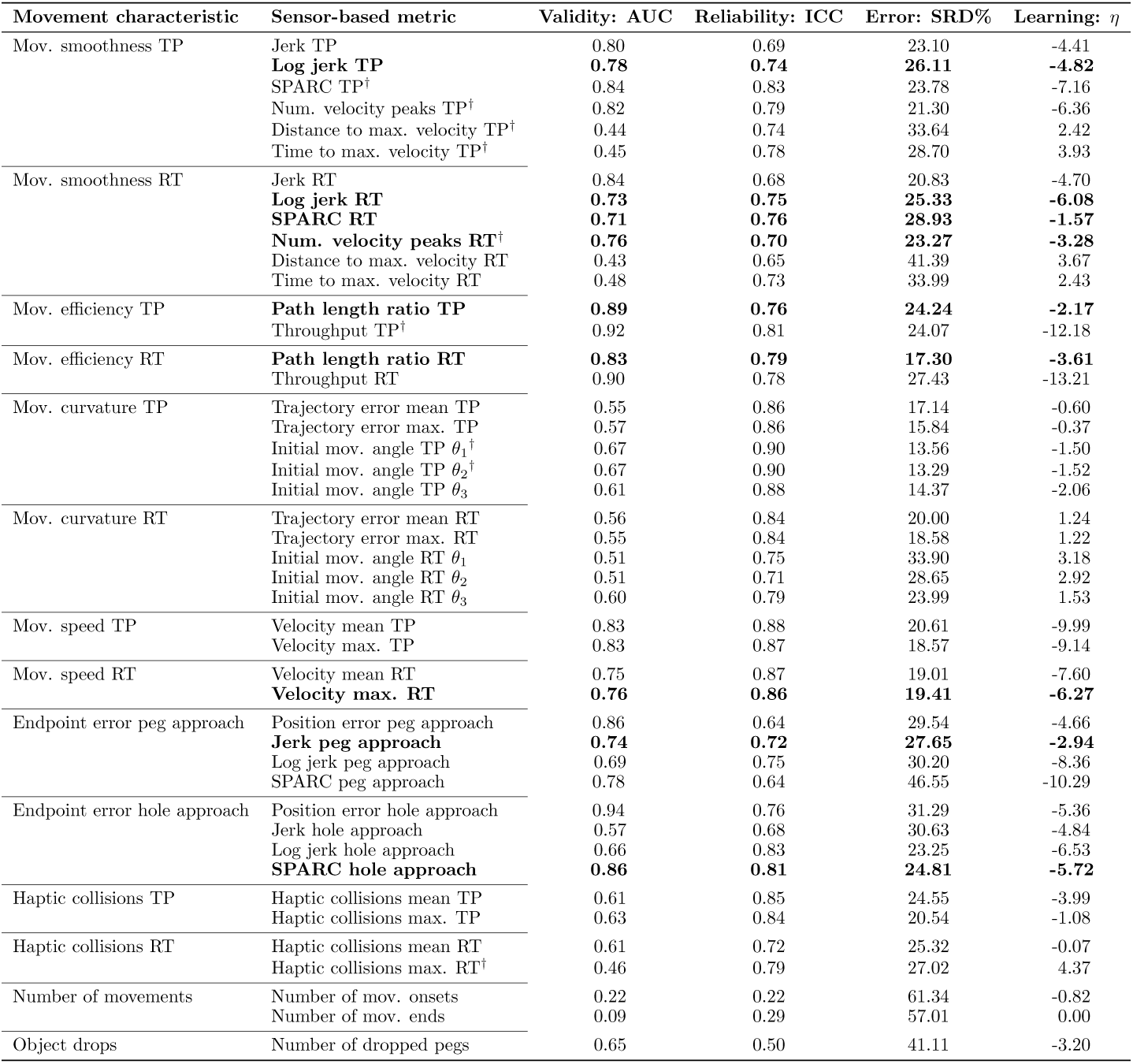
Results for the data-driven selection of kinematic metrics. The area under the curve (AUC, optimum at 1), intraclass correlation coefficient (ICC, optimum at 1), the smallest real difference (SRD%, optimum at 0), and *η* value (optimum at 0, worst at inf) were used to describe discriminative validity, test-retest reliability, measurement error, and learning effects, respectively. Metrics in bold fulfilled all evaluation criteria (AUC*>*0.7, ICC*>*0.7, SRD%*<*30.3, and *η >*-6.35). Metrics with insufficient model quality according to selection step 1 are annotated with a † and reported for completeness. mov: movement; TP: transport; RT: return; SPARC: spectral arc length; num: number.

| Movement characteristic | Sensor-based metric | Validity: AUC | Reliability: ICC | Error: SRD% | Learning: $\eta$ |
| --- | --- | --- | --- | --- | --- |
| Mov. smoothness TP | Jerk TP | 0.80 | 0.69 | 23.10 | -4.41 |
|  | **Log jerk TP** | **0.78** | **0.74** | **26.11** | **-4.82** |
|  | SPARC TP† | 0.84 | 0.83 | 23.78 | -7.16 |
|  | Num. velocity peaks TP† | 0.82 | 0.79 | 21.30 | -6.36 |
|  | Distance to max. velocity TP† | 0.44 | 0.74 | 33.64 | 2.42 |
|  | Time to max. velocity TP† | 0.45 | 0.78 | 28.70 | 3.93 |
| Mov. smoothness RT | Jerk RT | 0.84 | 0.68 | 20.83 | -4.70 |
|  | **Log jerk RT** | **0.73** | **0.75** | **25.33** | **-6.08** |
|  | **SPARC RT** | **0.71** | **0.76** | **28.93** | **-1.57** |
|  | **Num. velocity peaks RT†** | **0.76** | **0.70** | **23.27** | **-3.28** |
|  | Distance to max. velocity RT | 0.43 | 0.65 | 41.39 | 3.67 |
|  | Time to max. velocity RT | 0.48 | 0.73 | 33.99 | 2.43 |
| Mov. efficiency TP | **Path length ratio TP** | **0.89** | **0.76** | **24.24** | **-2.17** |
|  | Throughput TP† | 0.92 | 0.81 | 24.07 | -12.18 |
| Mov. efficiency RT | **Path length ratio RT** | **0.83** | **0.79** | **17.30** | **-3.61** |
|  | Throughput RT | 0.90 | 0.78 | 27.43 | -13.21 |
| Mov. curvature TP | Trajectory error mean TP | 0.55 | 0.86 | 17.14 | -0.60 |
|  | Trajectory error max. TP | 0.57 | 0.86 | 15.84 | -0.37 |
| | Initial mov. angle TP $\theta_1^\dagger$ | 0.67 | 0.90 | 13.56 | -1.50 |
| | Initial mov. angle TP $\theta_2^\dagger$ | 0.67 | 0.90 | 13.29 | -1.52 |
| | Initial mov. angle TP $\theta_3$ | 0.61 | 0.88 | 14.37 | -2.06 |
| Mov. curvature RT | Trajectory error mean RT | 0.56 | 0.84 | 20.00 | 1.24 |
|  | Trajectory error max. RT | 0.55 | 0.84 | 18.58 | 1.22 |
| | Initial mov. angle RT $\theta_1$ | 0.51 | 0.75 | 33.90 | 3.18 |
| | Initial mov. angle RT $\theta_2$ | 0.51 | 0.71 | 28.65 | 2.92 |
| | Initial mov. angle RT $\theta_3$ | 0.60 | 0.79 | 23.99 | 1.53 |
| Mov. speed TP | Velocity mean TP | 0.83 | 0.88 | 20.61 | -9.99 |
|  | Velocity max. TP | 0.83 | 0.87 | 18.57 | -9.14 |
| Mov. speed RT | Velocity mean RT | 0.75 | 0.87 | 19.01 | -7.60 |
|  | **Velocity max. RT** | **0.76** | **0.86** | **19.41** | **-6.27** |
| Endpoint error peg approach | Position error peg approach | 0.86 | 0.64 | 29.54 | -4.66 |
|  | **Jerk peg approach** | **0.74** | **0.72** | **27.65** | **-2.94** |
|  | Log jerk peg approach | 0.69 | 0.75 | 30.20 | -8.36 |
|  | SPARC peg approach | 0.78 | 0.64 | 46.55 | -10.29 |
| Endpoint error hole approach | Position error hole approach | 0.94 | 0.76 | 31.29 | -5.36 |
|  | Jerk hole approach | 0.57 | 0.68 | 30.63 | -4.84 |
|  | Log jerk hole approach | 0.66 | 0.83 | 23.25 | -6.53 |
|  | **SPARC hole approach** | **0.86** | **0.81** | **24.81** | **-5.72** |
| Haptic collisions TP | Haptic collisions mean TP | 0.61 | 0.85 | 24.55 | -3.99 |
|  | Haptic collisions max. TP | 0.63 | 0.84 | 20.54 | -1.08 |
| Haptic collisions RT | Haptic collisions mean RT | 0.61 | 0.72 | 25.32 | -0.07 |
|  | Haptic collisions max. RT† | 0.46 | 0.79 | 27.02 | 4.37 |
| Number of movements | Number of mov. onsets | 0.22 | 0.22 | 61.34 | -0.82 |
|  | Number of mov. ends | 0.09 | 0.29 | 57.01 | 0.00 |
| Object drops | Number of dropped pegs | 0.65 | 0.50 | 41.11 | -3.20 |

#### 2.4.1 Movement smoothness

Goal-directed movements are executed by translating parameters such as target distance into neural commands of certain amplitude, which are transferred to peripheral muscles performing a movement [50]. The signals’ amplitudes are chosen to minimize movement endpoint variance, which leads to smooth behaviour (i.e., bell-shaped velocity trajectories) [51]. These velocity trajectories can be modeled using a superposition of submovements and minimize the magnitude of the jerk trajectory [52]. In neurological subjects, more submovements with increased temporal shift and higher jerk magnitudes have been observed [53, 54], potentially due to disrupted feedforward control mechanisms. The temporal shift between subcomponents and the jerk magnitude was shown to reduce after receiving rehabilitation therapy [53], thereby highlighting their relevance to track recovery. We used the integrated jerk (referred to as *jerk*) normalized with respect to movement duration and length leading to a dimensionless metric to represent the intrinsic minimization of jerk [53]. The same metric was used with an additionally applied log transformation (*log jerk*) [55]. Additionally, the *spectral arc length* (i.e., metric describing spectral energy content) of the velocity trajectory should reflect the energy induced by jerky movements [55, 56]. Further, the number of peaks in the velocity profile (*number of velocity peaks*; MATLAB function *findpeaks*) was established as an inidicator for the number of submovements. Lastly, we calculated the time (*time to max. velocity*) and distance (*distance to max. velocity*) covered at peak velocity normalized with respect to the totally covered distance and time, respectively, to captures deviation from the typically observed bell-shaped velocity profile [57]. We calculated these metrics separately for *transport* and *return* as the *transport* requires precise grip force control, which could further affect feedforward control mechanisms.

#### 2.4.2 Movement efficiency

Ballistic movements in healthy subjects tend to follow a trajectory similar to the shortest path between start and target [58]. Previous studies suggested that neurologically affected subjects instead perform movements less close to the optimal trajectory compared to healthy controls [59] and that this behaviour correlates with impairment severity, as measured by the FMA-UE [60]. This suboptimal movement efficiency results in general from abnormal sensorimotor control, for example due to from erroneous state estimates for feedforward control, abnormal muscle synergy patterns (e.g., during shoulder flexion and abduction), weakness, and missing proprioceptive cues [47, 59, 61]. We used the path length ratio (i.e., shortest possible distance divided by the actually covered distance) to represent inefficient movements [59]. Additionally, the *throughput* (ratio of target distance and target width divided by movement time) was used as an information theory-driven descriptor of movement efficiency [62, 63]. The metrics were extracted from the start of the *transport* phase until the current peg was released and from the start of the *return* phase until the next peg was taken, as not only ballistic movements but also the endpoint error is of interest when describing the efficiency of movements.

#### 2.4.3 Movement curvature

While movement efficiency describes the overall deviation from the shortest path, it does not account for the direction of the spatial deviation. This might, however, be relevant to better discriminate abnormal feedfoward control from flexor synergy pattern or weakness, as in the latter two cases the movements might be especially performed closer to the body. We therefore selected five additional metrics to analyze the spatial deviation from the optimal trajectory in the horizontal plane [31, 32]. The *initial movement angle* was defined as the angular deviation between the actual and optimal trajectory [61]. As this metric requires the definition of a specific timepoint in the trajectory to measure the deviation, and as multiple approaches were used in literature [57, 61, 62, 64], we explored three different ways to define the timepoint. This included the time at which 20% of the shortest distance between peg and hole was covered (*initial movement angle θ*_1_), the time at which 20% of the actually covered distance between peg and hole was reached *initial movement angle θ*_2_, and the time at which peak velocity was achieved (*initial movement angle θ*_3_). Additionally, the *mean* and *maximal trajectory error* with respect to the ideal, straight trajectory were calculated. All metrics were estimated separately for *transport* and *return*.

#### 2.4.4 Movement speed

The speed of ballistic movements in healthy subjects is mostly controlled by the tradeoff between speed and accuracy as described by Fitt’s law, which is indirectly imposed through the concept of velocity-dependent neural noise [51, 63]. In neurologically affected subjects, increased speed can, for example, result from inappropriately scaled motor commands and disrupted feedforward control [47]. On the other hand, reduced speed can also stem from weakness (i.e., reduced ability to active spinal motor neurons leading to decreased strength) or spasticity (i.e., velocity-dependent increase in muscle tone), the latter resulting from upper motor neuron lesions, abnormally modulated activity in the supraspinal pathways, and thereby increased hyperexcitability of stretch reflexes [47, 65]. We calculated the mean (*velocity mean*) and maximum (*velocity max.*) values of the velocity trajectory to represent movement speed during the *transport* and *return* phases.

#### 2.4.5 Endpoint error

To fully characterize the speed-accuracy tradeoff, we additionally analyzed the position error at the end of a movement. In neurological disorders, increased endpoint error (i.e., dysmetria) was commonly observed and can, for example, result from inappropriately scaled motor commands and thereby disrupted feedforward control [66, 67], but also from cognitive and proprioceptive deficits [68]. Dysmetria was found especially in post-stroke subjects with lateral-posterior thalamic lesions [68], is a common manifestation of intention tremor in MS [69], and is typically observed in subjects with cerebellar ataxia [70]. In the VPIT, the cumulative horizontal Euclidean distance between the cursor position and targeted peg or hole (*position error*) were calculated during the *peg approach* and *hole approach* phases, respectively. Further, the *jerk*, *log jerk*, and *spectral arc length* metrics were calculated during both phases, as a jerk index was shown previously to correlate with the severity of intention tremor in MS [71].

#### 2.4.6 Haptic collisions

Haptic collisions describe the interaction forces between a subject and the virtual pegboard rendered through the haptic device. Haptic guidance can be used to ease inserting the virtual pegs into the holes, which have reduced haptic impedance. Previous studies indicated increased haptic collision forces in multiple neurological disorders and especially stroke subjects with sensory deficits [29, 72]. We additionally expected that collision forces during *transport* and *return* (i.e., phases during which haptic guidance is not required) could be increased due to arm weakness. In particular, neurological subjects can have a limited capability to lift their arm against gravity, leading to increased vertical haptic collisions [73]. The mean and max. vertical collision force (*haptic collisions mean* and *haptic collisions max.*) was calculated during *transport* and *return* to quantify haptic collision behaviour.

#### 2.4.7 Number of successful movements

Subjects without neurological deficits can start and end goal-directed movements with ease. On the contrary, persons with neurological disorders can have a reduced ability to initiate and terminate ballistic movements with potentially heterogeneous underlying impairments including abnormal feedforward control, sensory feedback, spasticity, weakness, and fatigue [13, 47, 57]. Therefore, the metric *number of movement onsets* was defined based on the number of valid pegs, using the defined segmentation algorithm, when identifying the start of the *transport* and *return* phases. Analogously, *number of movement ends* was based on the sum of correctly segmented ends for the *transport* and *return* phases.

#### 2.4.8 Object drops

Neurologically intact subjects can precisely coordinate arm movements and finger forces to transport objects. This ability can be reduced in neurological disorders and can potentially lead to the drop of an object during its transport [74]. Underlying mechanisms include for example distorted force control due to incorrectly scaled motor commands or distorted sensory feedback as well as reduced spatio-temporal coordination between arm and hand movements [47, 74]. In the VPIT, the number of virtual pegs that were dropped (*dropped pegs*) should represent object drops and thereby grip force control as well as the spatio-temporal coordination of arm and hand movements. The metric was defined based on how often the grasping force dropped below a 2 N threshold (i.e., subjects still holding the handle) while lifting a virtual peg [32].

#### 2.4.9 Grip force scaling and coordination

The precise scaling and spatio-temporal coordination of grasping forces is a key requirement for successful object manipulation and leads, in neurologically intact subjects, to single-peaked bell-shaped grip force rate profiles when starting to grasp objects [75]. Abnormal grip force scaling and decreased grip force coordination have been reported in neurological subjects, resulting in multi-peaked grip force rate profiles, and were attributed to, for example, distorted feedforward control, abnormal somatosensory feedback and processing, as well as the presence of the pathological flexor synergy [75–82]. Also, a reduction in applied grip force levels due to weakness can be expected depending on the neurological profile of a subject [47]. Further, a slowness of *force buildup* [77] and *force release* [78] has been reported, even though other studies showed that the ability to produce and maintain submaximal grip forces was preserved [74, 78]. Additionally, there is evidence suggesting that *force buildup* and *force release* have different neural mechanisms and that force control can further be decomposed into force scaling and motor coordination [78, 79].

To describe grip force scaling, we applied four metrics separately to the *transport*, *return*, *peg approach*, and *hole approach* phases. We calculated the mean (*grip force mean*) and maximum (*grip force max.*) value of the grasping force signal during each phase. Additionally, we estimated the mean absolute value (*grip force rate mean*) and absolute maximum (*grip force rate max.*) of the grip force rate time-series. Similarly, we characterized grip force coordination during the *transport*, *return*, *peg approach*, *hole approach*, *force buildup* and *force release* phases, for which we calculated the number of positive and negative extrema (*grip force rate number of peaks*) and the spectral arc length (*grip force rate spectral arc length*). For the *force buildup* and *force release* phases, which contain only the segments of most rapid force generation and release, respectively, we additionally calculated their duration (*force buildup/release duration*).

#### 2.4.10 Overall disability

A single indicator expected to describe the subject-specific *overall disability* level was defined based on the *task completion time* (i.e., duration from first *transport* phase until insertion of last peg).

### 2.5 Data postprocessing

To reduce the influence of intra-subject variability, the grand median across pegs and repetitions was computed for each metric. Subsequently, the influence of possible confounds, which emerge from subject demographics not related to neurological disorders, was modeled based on data from all neurologically intact subjects. This should allow to compensate for these factors when analyzing data from neurologically affected subjects. In more detail, the impact of age (in yrs), sex (male or female), tested body side (left or right), and handedness (performing the test with the dominant side: true or false) were used as fixed effects (i.e., one model slope parameter per independent variable) in a linear mixed effect model generated for each sensor-based metric [83]. Additionally, the presence of stereo vision deficits (true or false) was used as a fixed effect, as the perception of depth in the VR environments might influence task performance [84, 85]. A subject-specific random effect (i.e., one model intercept parameter per subject) was added to account for intra-subject correlations arising from including both tested body sides for each subject. A Box-Cox transformation was applied on each metric to correct for heteroscedasticity, as subjectively perceived through non-normally distributed model residuals in quantile-quantile plots [86]. Additionally, this transformation allows to capture non-linear effects with the linear models. The models were fitted using maximum likelihood estimation (MATLAB function *fitlme*) and defined as:

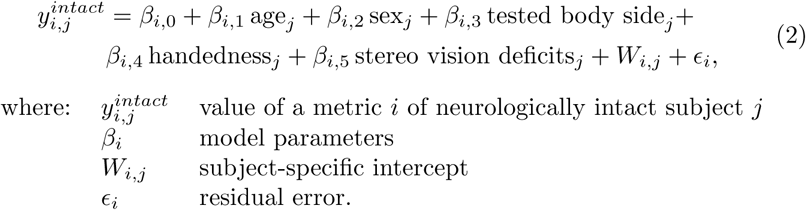

For any subject being analyzed, the effect of all confounds on the sensor-based metric was removed based on the fitted models. This generated the value *ӯ_i,j_* of a metric without confounds arising from subject demographics:

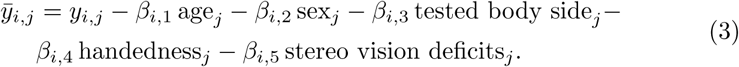

Furthermore, the corrected values *ӯ_i,j_* were then expressed relative to all neurologically intact subjects (*ӯ_i_^intact^*) with the goal to standardize the range of all metrics, which simplifies their physiological interpretation and enables the direct comparison of different metrics. Therefore, the normalized value *ŷ_i,j_* was defined relative to the median and variability *d_i_* of all neurologically intact subjects:

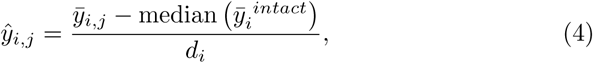

with the median absolute deviation (MAD) of all neurologically intact subjects being used as a variability measure [87]:

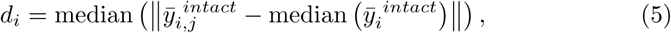

The MAD was preferred over the standard deviation, as the former allows a more robust analysis that is independent of the underlying distribution of a metric [87]. Lastly, the values *ŷ_i,j_* were divided by the maximal observed value in the included neurological population, such that the subject currently showing worst task-performance receives a score of 100%. In order to discriminate normal from abnormal behaviour based on the normalized values, a cut-off was defined based on the 95*^th^* percentile (i.e., imposed false positive detection rate of 5%) of each metric *ŷ*^*i*^^*intact*^ across all neurologically intact subjects.

### 2.6 Data-driven selection and validation of sensor-based metrics

The sensor-based metrics were reduced to a subset with optimal clinimetric properties based on three selection steps, followed by two additional validation steps. To evaluate the ability of this selection process to discriminate between physiologically-relevant information and random noise, the selection steps were additionally applied to a simulated random metric (*simulated Gaussian noise*) containing no physiologically relevant information. This metric was constructed by randomly drawing data from a log-normal distribution (mean 46.0, standard deviation 32.2, mimicking the distribution of the *total time* for the *reference population*) for each subject and tested body side.

#### Metric selection & validation: step 1

With the goal to better understand the influence of subject demographics on the sensor-based metric, simulated likelihood ratio tests (1000 iterations) between the full model and a reduced model without the fixed effect of interest were used to generate *p*-values that were interpreted based on a 5% significance level [88]. This allowed to judge whether a fixed effect influenced the sensor-based metric in a statistically significant manner. We removed metrics that were significantly influenced by stereo vision deficits, as we expected that the influence of stereo vision deficits can not always be compensated for, for example if their presence is not screened in a clinical setting.

As the performance of the presented confound correction process depends on the fit of the model to the data, we additionally removed metrics with low model quality according to the criteria *C*1 and *C*2, which describe the mean absolute estimation error (MAE) of the models and its variability [89]:

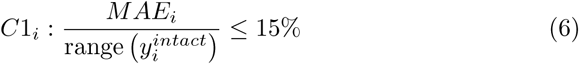

and

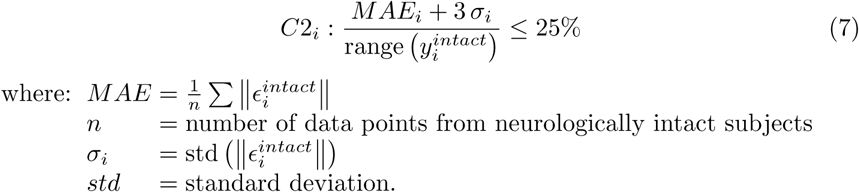

Fulfilling both criteria leads to the selection of models with *moderate* and *good* quality according to the definition of Roy et al. [89]. Before the calculation of C1 and C2, data points with the 5% highest residuals were removed [89]. The criteria C1 and C2 were preferred over the more commonly used coefficient of determination *R*^2^, because the magnitude of this metric is highly dependent on the distribution of the dependent variable, which prohibits the definition of a model quality threshold that is valid across metrics [89, 90].

#### Metric selection & validation: step 2

Receiver operating characteristic (ROC) analysis was used to judge the potential of a metric to discriminate between neurologically intact and affected subjects, which is a fundamental requirement to validate that the proposed metrics are sensitive to sensorimotor impairments [22, 91]. In more detail, a threshold was applied for each metric to classify subjects as being either neurologically intact or impaired. The threshold was varied across the range of all observed values for each metric and the true positive rate (number of subjects correctly classified as neurologically affected divided by the total number of neurologically affected subjects) and false positive rate (number of subjects incorrectly classified as neurologically affected divided by the total number of neurologically intact subjects) were calculated. The area under the curve (AUC) when plotting true positive rates against false positive rates was used as a quality criterion for each metric (Figure 2).

**Figure 2:**
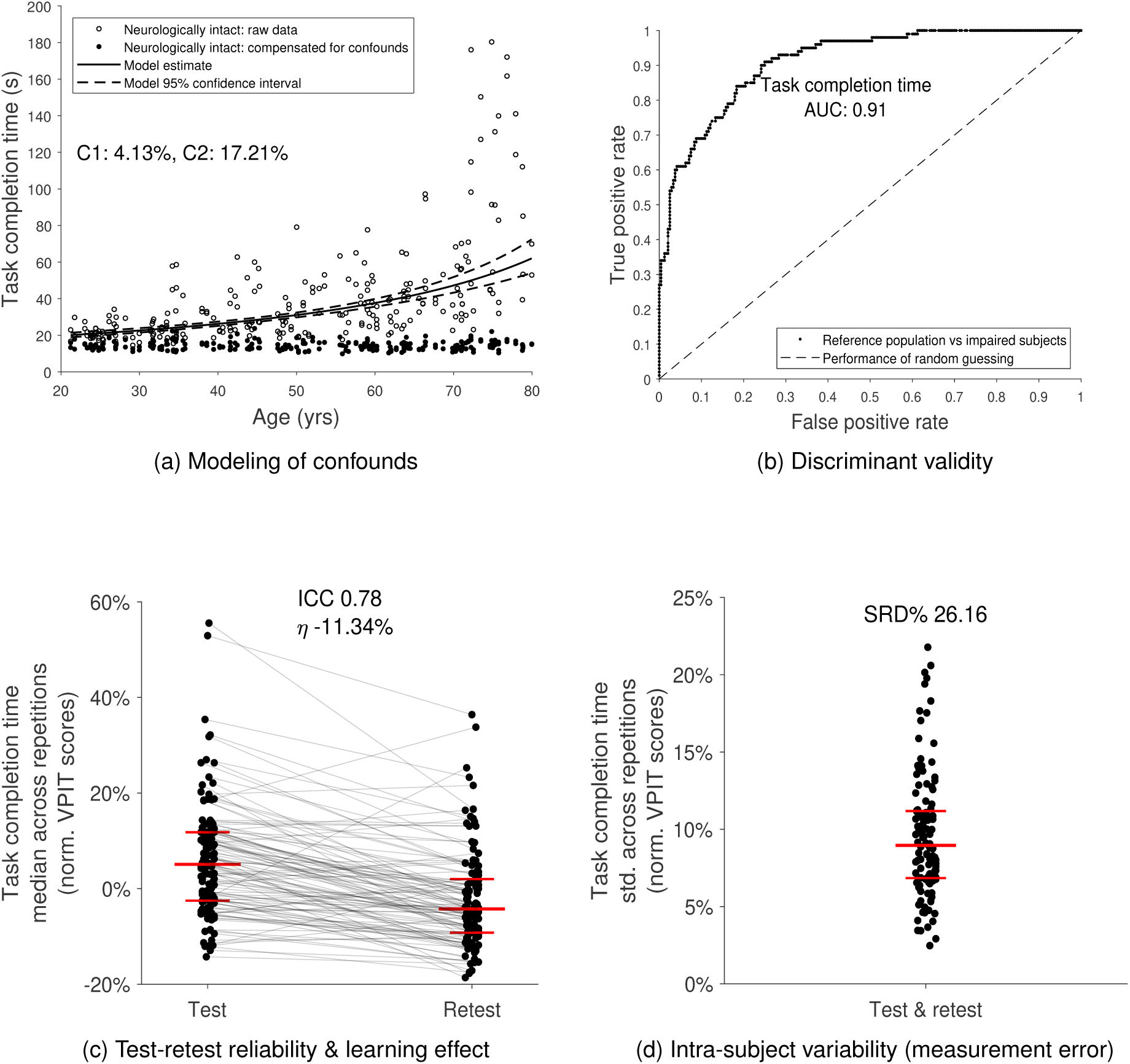
Data-driven selection and validation of metrics: example of task completion time. **(a)** the influence of age, sex, tested body side, handedness, and stereo vision deficits on each digital health metrics was removed using data from neurologically intact subjects and mixed effect models (model quality criteria *C*1 and *C*2). Models were fitted in a Box-Cox-transformed space and back-transformed for visualization. Metrics with low model quality (*C*1 *>*15% or *C*2 *>*25%) were removed. **(b)** The ability of a metric to discriminate between neurologically intact and affected subjects (discriminant validity) was evaluated using the area under the curve value (AUC). Metrics with AUC*<*0.7 were removed. **(c)** Test-retest reliability was evaluated using the intra-class correlation coefficient (ICC) indicating the ability of a metric to discriminate between subjects across testing days. Metrics with ICC*<*0.7 were removed. Additionally, metrics with strong learning effects (*η >*-6.35) were removed. The long horizontal red line indicates the median, whereas the short ones represent the 25*^th^*- and 75*^th^*-percentile. **(d)** Measurement error was defined using the smallest real difference (SRD%), indicating a range of values for that the assessment cannot discriminate between measurement error and physiological changes. The distribution of the intra-subject variability was visualized, as it strongly influences the SRD. Metrics with SRD%*>*30.3 were removed.

For metrics to be responsive to intervention-induced physiological changes and allow a meaningful tracking of longitudinal changes, it is fundamental to have low intra-subject variability, high inter-subject variability, and yield repeatable values across a test-retest sessions. Therefore, the data set with 60 neurologically intact subjects performing the VPIT protocol on two separate testing days was used to quantify test-retest reliability. Specifically, the intra-class correlation coefficient (ICC) was calculated to describe the ability of a metric to discriminate between subjects across multiple testing days (i.e., inter-subject variability) [92, 93]. The agreement ICC based on a two-way analysis of variance (ICC A,k) was applied while pooling data across both tested body sides. Further, the smallest real difference (SRD) was used to define a range of values for that the assessment cannot distinguish between measurement error and an actual change in the underlying physiological construct (i.e., intra-subject variability) [94]. For each metric *i*, the SRD was defined as

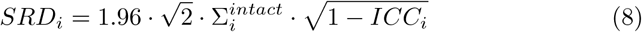

where: Σ*_i_* = std across repetitions, subjects, and testing days.

To directly relate the SRD to the distribution of a metric, it was further expressed relative to a metrics’ range:

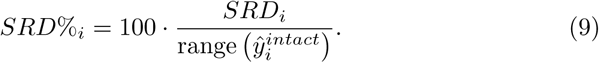

Lastly, to distinguish task-related learning from physiological changes when testing subjects before and after receiving an intervention, the presence and strength of learning effects was calculated for each metric. For this purpose, a paired *t*-test was performed between data collected at test- and retest to check for a statistically significant difference between the days. Then, the strength (i.e., slope) of the learning effect was estimated by calculating the mean difference between test and retest and normalizing it with respect to the range of observed values:

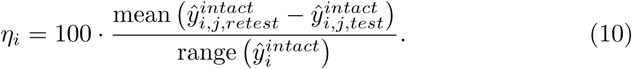

Metrics passed this second selection step if the AUC did indicate acceptable, excellent, or outstanding discriminant ability (AUC ≥ 0.7) and they had at least acceptable reliability (i.e., ICC values above 0.7) [22, 91]. As no cutoff has been defined for the interpretation of the SRD% [95], we removed the metrics that had the 20% worst SRD% values. Hence, metric passed the evaluation (i.e., small measurement error relative to other metrics) if the SRD% was below 30.3 (80*^th^*-percentile). Similarly, no cutoff for the interpretation of learning effects was available. Hence, metrics passed the evaluation (i.e., no strong learning effects) if *η* was above −6.35 (20*^th^*-percentile) of observed values.

#### Metric selection & validation: step 3

The correlations between the metrics were analyzed with the goal to identify a set of metrics that contains little redundant information to simplify clinical interpretability. Therefore, a correlation matrix was constructed using partial Spearman correlations. This technique allows to describe the relation between two metrics and to simultaneously model all other metrics that could potentially influence the relationship between the two metrics of interest [96, 97]. Hence, this approach can help to exclude certain non-causal correlations. A pair of metrics with an absolute partial correlation *ρ_p_* of at least 0.5 was considered for removal [98]. From this pair of metric, the one that had inferior psychometric properties (AUC, ICC, and SRD%) or was less accepted in literature was removed. To simplify the interpretation of the correlation results, we applied the analysis only to metrics that passed all previous selection steps. Additionally, this analysis was applied in an iterative manner, as the removal of certain metrics, which were previously modeled, can change the remaining inter-correlations. The correlation coefficients were interpreted according to Hinkle et al.: very high: *ρ_p_ ≥*0.9; high: 0.7*≤ ρ_p_ <*0.9; moderate: 0.5*≤ ρ_p_ <*0.7; low: 0.3*≤ ρ_p_ <*0.5; very low: *ρ_p_ <*0.3 [98].

#### Further validation of metrics: step 1

To better identify the pathophysiological correlates of the metrics that passed all previous evaluation steps, exploratory factor analysis was applied [99–101]. This method tries to associate the variability observed in all metrics with *k* unobserved latent variables via factor loadings, which can be interpreted in light of the initial physiological motivation of the metrics. Exploratory factor analysis was implemented using maximum likelihood common factor analysis followed by a *promax* rotation (MATLAB function *factoran*). For the interpretation of the emerged latent space, we only considered strong (absolute value≥0.5) factor loadings [99]. The number of factors *k* was estimated in a data-driven manner using parallel analysis (R function *fa.parallel*) [102]. This approach simulates a lower bound that needs to be fulfilled by the eigenvalue associated to each factor and has been shown to be advantageous compared to other more commonly used criteria, such as the Kaiser condition (i.e., eigenvalues*>*1 are retained) [100, 101]. Also, the Kaiser-Meyer-Olkin value (KMO) was calculated to evaluate whether the data was mathematically suitable for the factor analysis.

#### Further validation of metrics: step 2

An additional clinically-relevant validation step evaluated the ability of the metrics to capture the severity of upper limb disability. For this purpose, each population was grouped according to their disability level as defined by commonly used clinical scores. Subsequently, the behaviour of the metrics across the sub-populations and the *reference population* were statistically analyzed. Stroke subjects were grouped according to the FMA-UE score (ceiling: FMA-UE=66; mild impairment: 54*≤* FMA-UE*<*66; moderate impairment: 35*≤* FMA-UE*<*54) [103]. MS subjects were split into three groups based on their ARAT score (full capacity: 55*≤* ARAT*≤* 57; notable capacity: 43 *≤*ARAT*<*55; limited capacity: 22 *≤*ARAT*<*43) [104]. ARSACS subjects were divided into three different age-groups (young: 26*≤* age*≤* 36; mid-age: 37 *≤*age *≤*47; older-age: 48*≤*age*≤*58) due to the neurodegenerative nature of the disease [4]. A Kruskal-Wallis omnibus test followed by post-hoc tests (MATLAB functions *kruskalwallis* and *multcompare*) were applied to check for statistically significant differences between groups. Bonferroni corrections were applied in both cases.

## 3 Results

Data from 120 neurologically intact subjects of age 51.1 [34.6, 65.6] yrs (median [25*^th^*-percentile, 75*^th^*-percentile]; 60 male; 107 right hand dominant; 12 with stereo vision deficits) was acquired with 60 of them performing a test-retest session (age 48.8 [40.2, 60.2]; 34 male; 48 right hand dominant; time between sessions 5.0 [4.0, 6.5] days). Eighty-nine neurologically affected subjects (53 post-stroke with affected side FMA-UE 57 [49, 65], 28 MS with ARAT 52.0 [46.5, 56.0], 8 ARSACS with NHPT 43.5 [33.1, 58.7] s) were used for the selection and validation of the metrics. Their age was 56.2 [42.1, 65.3] yrs, 52 were male, 75 were right hand dominant, and for 35 stroke subjects, the right body side was most affected. In total, data from 43350 individual movements were recorded. Detailed demographic and the available clinical information for each neurologically affected subject can be found in Table SM1.

### Selection of metrics: step 1

The influence of all potential confounds and the model quality for each sensor-based metric including *p*-values can be found in Table SM2 (example in Figure 2). For all metrics, 69.7%, 44.7%, 27.6%, 6.6%, and 7.9% were significantly influenced by age, sex, tested side, hand dominance, and stereo vision deficits, respectively. In more detail, *initial movement angle transport θ*_1_*, θ*_2_*, θ*_3_, *number of movement ends*, *number of dropped pegs*, *grip force rate number of peaks buildup* were the metrics being altered by stereo vision deficits. The required quality of the models, according to the C1 and C2 criteria, were not fulfilled by thirteen (16.9%) of all metrics (including the *simulated Gaussian noise*, see SM for a detailed list).

### Selection of metrics: step 2

Thirteen (16.9%) out of 77 metrics fulfilled the criteria of the validity, reliability, measurement error, and learning analysis (Figure 2, Table 1, and Table 2). The median AUC, ICC, SRD%, and *η* values of the 12 metrics that passed step 1 and step 2 were 0.77 [0.74, 0.85], and 0.80 [0.75, 0.82], 24.6 [21.5, 26.2], and −5.72 [−6.09, −3.27] respectively. The *simulated Gaussian noise* metric did not pass this evaluation step (AUC 0.37, ICC −0.07, SRD% 117.04, *η* 0.25).

**Table 2:**
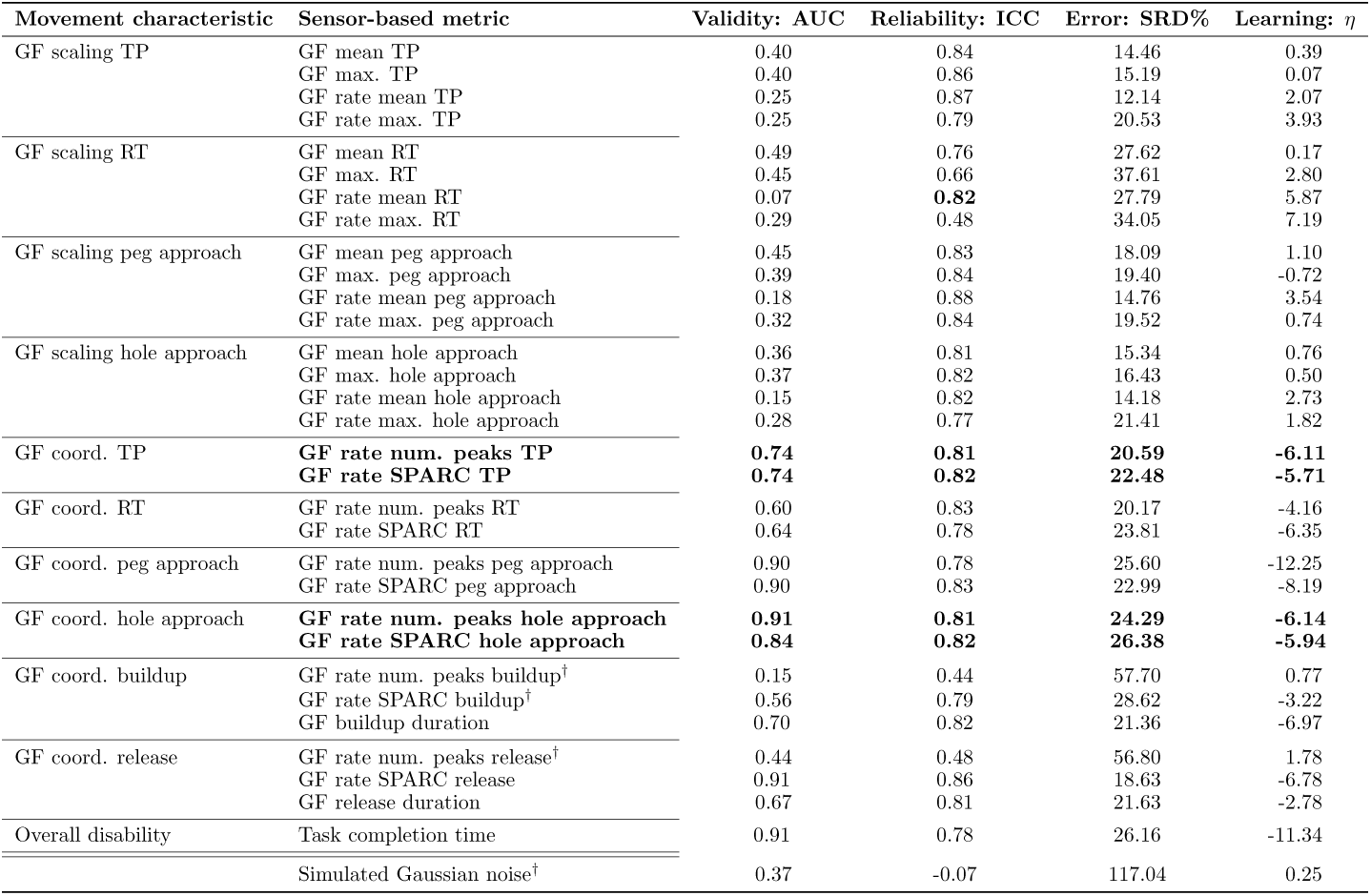
Results for the data-driven selection of kinetic metrics. The area under the curve (AUC, optimum at 1), intraclass correlation coefficient (ICC, optimum at 1), the smallest real difference (SRD%, optimum at 0), and *η* value (optimum at 0, worst at inf) were used to describe discriminative validity, test-retest reliability, measurement error, and learning effects, respectively. The *task completion time* and the *simulated Gaussian noise* metrics were evaluated in addition to the kinetic metrics. Rows in bold fulfilled all evaluation criteria (AUC*>*0.7, ICC*>*0.7, SRD%*<*30.3, and *η >*-6.35). Metrics with insufficient model quality according to selection step 1 are annotated with a † and reported for completeness. GF: grip force; TP: transport; RT: return; SPARC: spectral arc length; num: number.

### Selection of metrics: step 3

The constructed partial correlation matrices can be found in Figure 3. Among the remaining metrics, *grip force rate number of peaks hole approach* was removed as it correlated (*ρ_p_* ≥ 0.5) with *grip force rate spectral arc length approach hole* and the latter metric is less influenced through confounds as it is independent of movement distance. Additionally, *spectral arc length hole approach* was discarded as it correlated with *grip force rate spectral arc length hole approach* and the latter metric is more directly related to hand function, which was not yet well covered by the other metrics. The remaining 10 metrics yielded absolute partial inter-correlations of 0.14 [0.06 0.24] (zero very high, zero high, zero moderate, six low, and 39 very low inter-correlations).

**Figure 3:**
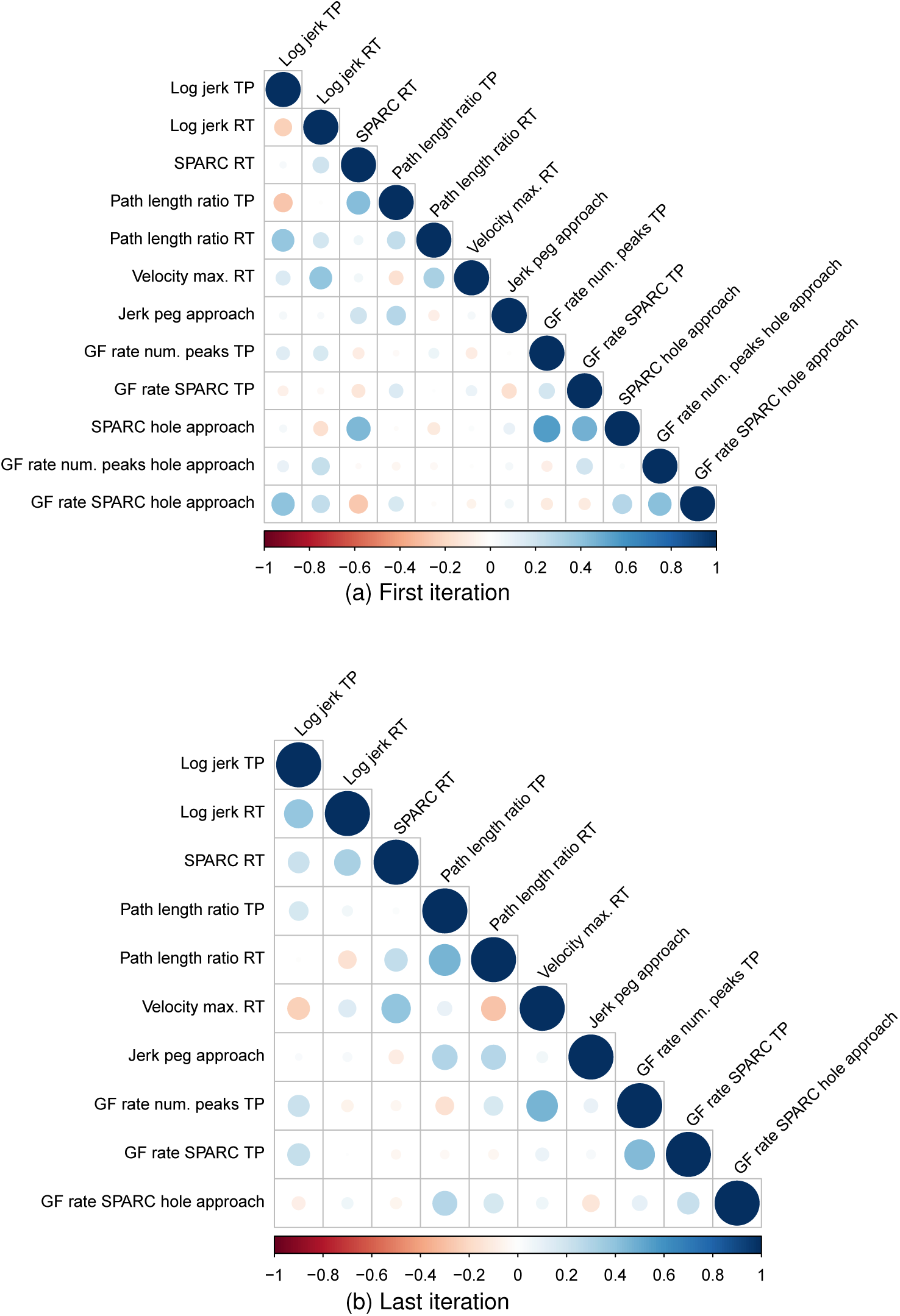
Partial correlation analysis. The objective was to remove redundant information. Therefore, partial Spearman correlations were calculated between all combination of metrics while controlling for the potential influence of all other metrics. Pairs of metrics were considered for removal if the correlation was equal or above 0.5 The process was done in an iterative manner and the first and the last iterations are presented.

### Further validation of metrics: step 1

The Kaiser-Meyer-Olkin value was 0.82, which indicated that the application of the factor analysis was suitable [105, 106]. According to the parallel analysis, the most likely number of underlying latent factors *k* was five (Figure SM2). The factor loadings can be found in Table 3. The metrics *path length ratio transport/return* and *jerk peg approach* had strong loadings on factor 1. The metrics *log jerk transport*, *log jerk return*, and *spectral arc length return* loaded strongly on factor 2. The metrics *grip force rate number of peaks transport* and *grip force rate spectral arc length transport* had strong loadings on factor 3, whereas *velocity max. return* and *grip force rate spectral arc length hole approach* loaded strongly on factor 4 and 5, respectively.

**Table 3:** Structural validity: exploratory factor analysis. Loadings of metrics on underlying latent factors extracted with exploratory factor analysis. The interpretation of each metric was physiologically motivated initially. Larger absolute loadings indicate a stronger contribution to a factor. Bold font indicate strong loadings (i.e., absolute loading of at least 0.5). Abbreviations: F1-5: data-driven latent factors. GF: grip force; coord: coordination; num: number; SPARC: spectral arc length.

| Expected interpretation | Sensor-based metric | F1 | F2 | F3 | F4 | F5 |
| --- | --- | --- | --- | --- | --- | --- |
| Movement smoothness transport | Log jerk transport | 0.09 | **0.73** | 0.21 | -0.19 | -0.05 |
| Movement smoothness return | Log jerk return | -0.08 | **0.86** | -0.11 | 0.02 | 0.02 |
|  | SPARC return | 0.10 | **0.59** | -0.10 | 0.23 | -0.03 |
| Movement efficiency transport | Path length ratio transport | **0.83** | 0.08 | -0.17 | 0.06 | 0.11 |
| Movement efficiency return | Path length ratio return | **0.79** | -0.06 | 0.08 | -0.14 | 0.04 |
| Movement speed transport | Velocity max. return | -0.02 | 0.01 | 0.16 | **0.90** | 0.01 |
| Endpoint error peg approach | Jerk peg approach | **0.72** | -0.04 | 0.12 | 0.07 | -0.14 |
| GF coord. transport | GF num. peaks transport | 0.00 | -0.06 | **0.93** | 0.11 | -0.03 |
|  | GF rate SPARC transport | -0.08 | 0.19 | **0.62** | 0.00 | 0.11 |
| GF coord. hole approach | GF rate SPARC hole approach | 0.11 | -0.02 | 0.02 | 0.01 | **0.94** |

### Further validation of metrics: step 2

The behaviour of all metrics across subject subpopulations with increasing disability level can be found in Figure 4, 5, and 6. All metrics indicated statistically significant differences between the neurologically intact and at least one of the neurologically affected subpopulations for each disorder, with the exception of *jerk peg approach* in MS subjects. Additionally, significant differences between subpopulations were found for *log jerk transport* in stroke subjects. Consistent trends (i.e., monotonically increasing medians across subpopulations) were found for all metrics except for *spectral arc length return*, *force rate spectral arc length approach hole*, and *force rate num. peaks approach hole*.

**Figure 4:**
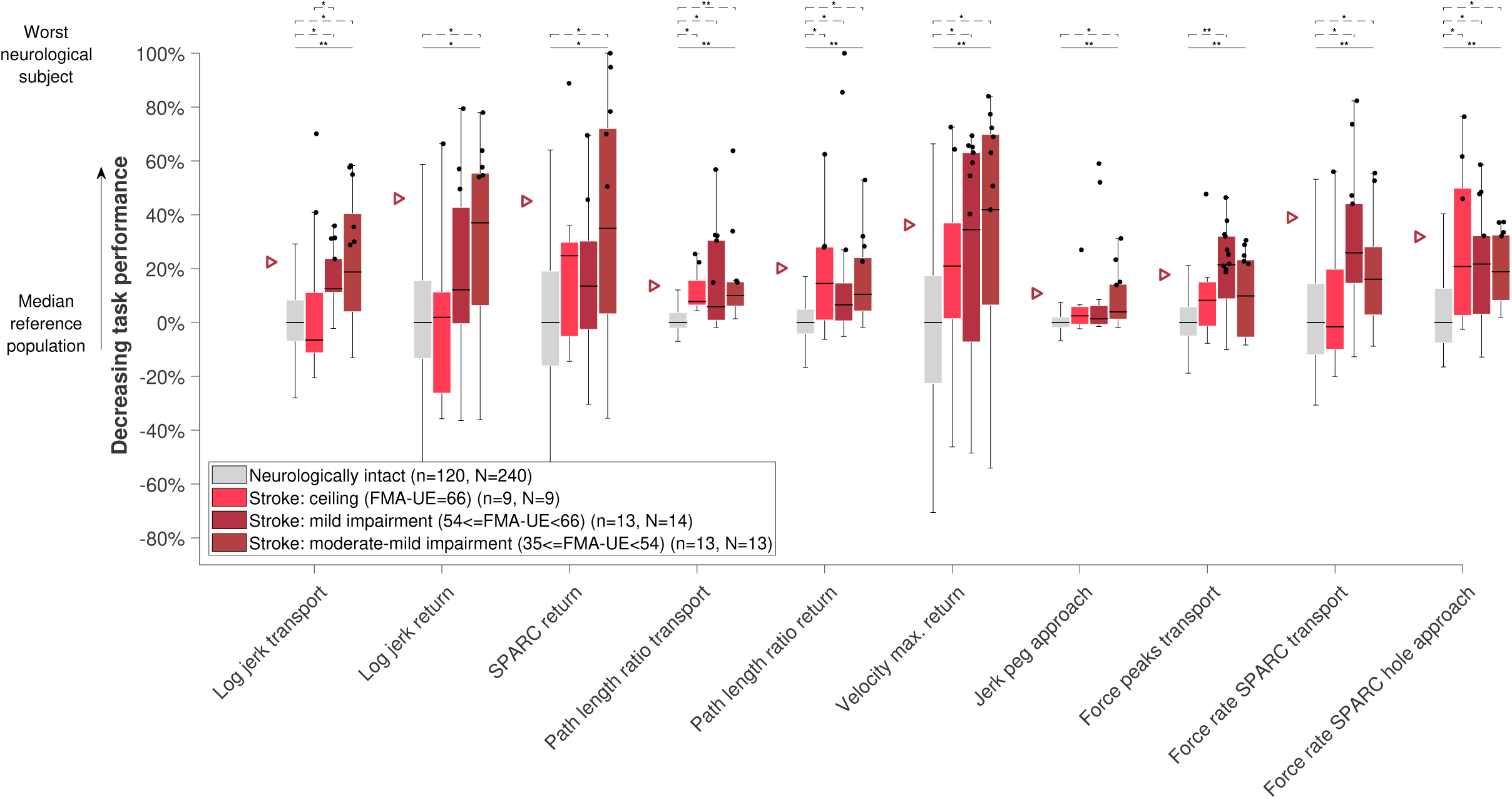
Sensitivity of metrics to disability severity in stroke subjects. The subject population was grouped according to their clinical disability level and compared between the subpopulations and to neurologically intact subjects. The vertical axis in the sensorimotor profile indicates task performance based on the distance to the reference population. In the box plots, the median is visualized through the black horizontal line, the interquartile range (IQR) through the boxes, and the minimum and maximum value within 1.5 IQR of the lower and upper quartile, respectively, through the whiskers. Single data points above the 95*^th^*-percentile (indicated with triangles) of neurologically intact subjects are defined as showing abnormal behaviour and are represented with black dots. Solid and dashed horizontal black lines above the box plots indicate results of the omnibus and post-hoc statistical tests, respectively. Only significant *p*-values after Bonferroni correction were visualized (*indicates *p <* 0.05 and \*\**p <* 0.001). The value *n* refers to the number of subjects in that group and *N* to the number of data points. Only subjects with available clinical scores were used for the analysis. For the *jerk peg approach*, one outlier data point was not visualized to maintain a meaningful representation. SPARC: spectral arc length.

**Figure 5:**
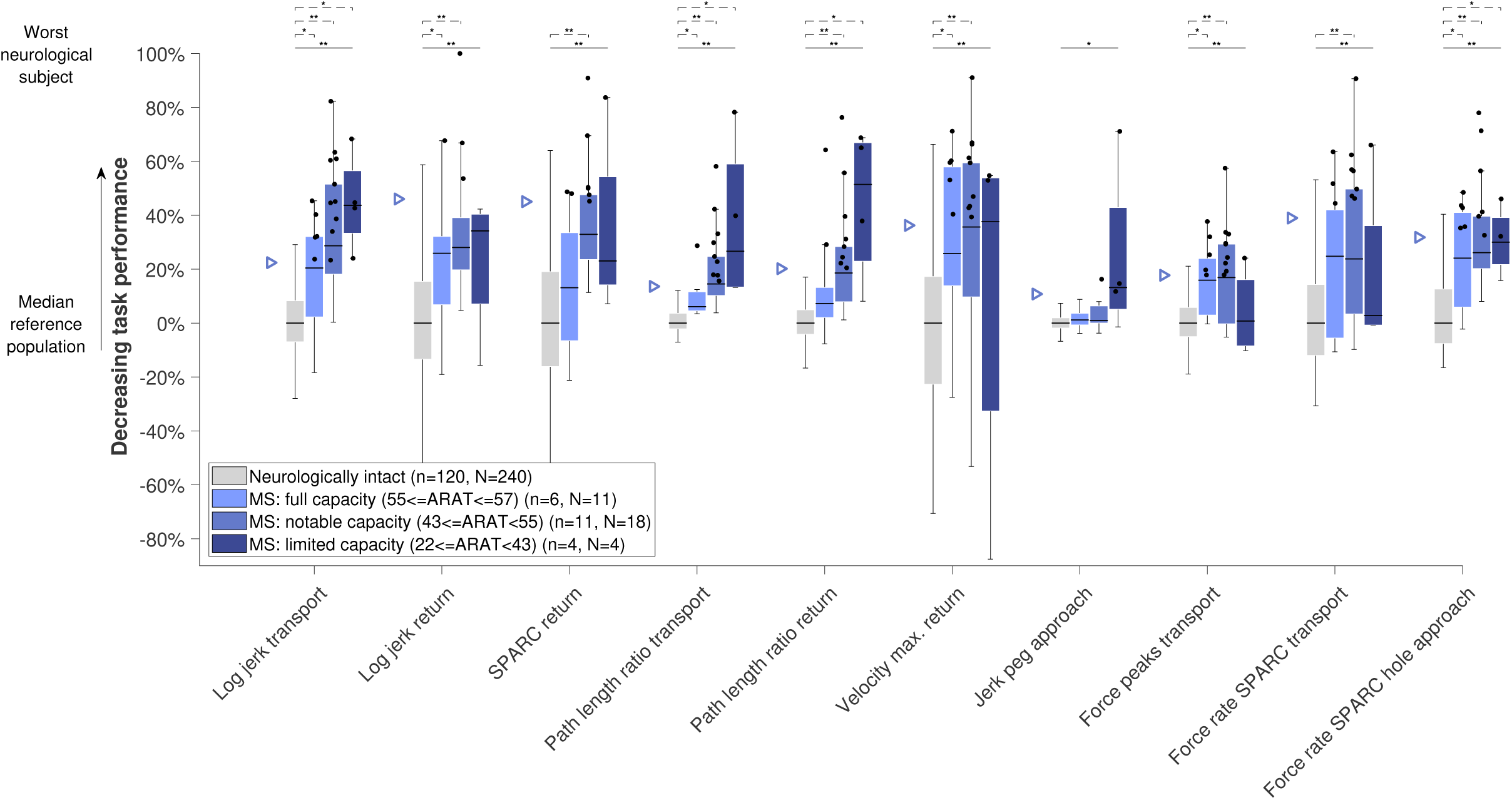
Sensitivity of metrics to disability severity in MS subjects. See Figure 4 for a detailed description.

**Figure 6:**
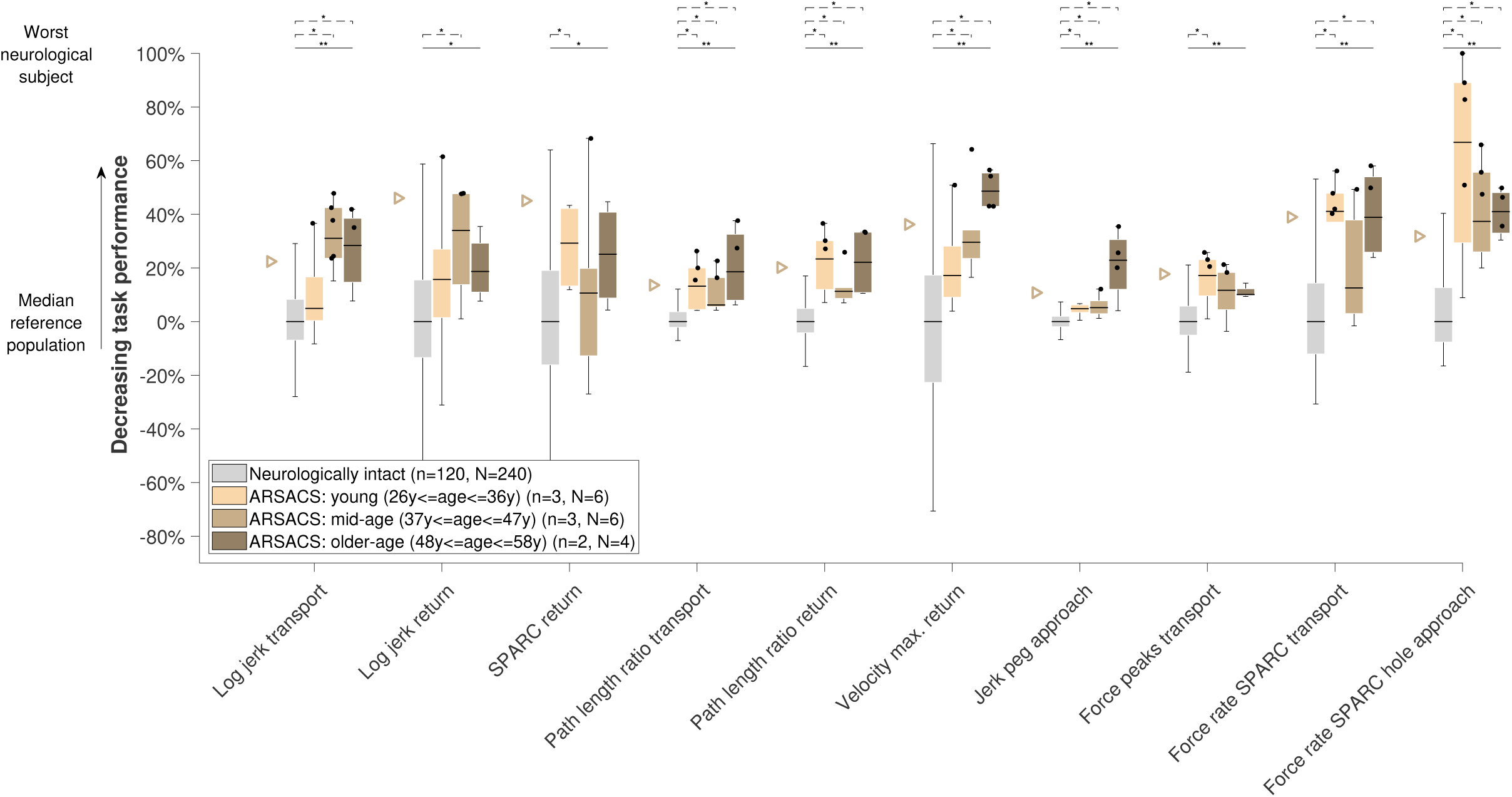
Sensitivity of metrics to disability severity in ARSACS subjects. See Figure 4 for a detailed description.

## 4 Discussion

In this work, we aimed to propose and apply a data-driven framework to select and validate digital health metrics, with the objective to facilitate their still lacking clinical integration. The approach considers i) the targeted impairments, ii) the influence of participant demographics, and iii) important clinimetric properties. As an example use-case, we implemented this framework with 77 kinematic and kinetic metrics extracted from the VPIT, a previously proposed sensor-based assessment of arm and hand sensorimotor impairments. For this purpose, the VPIT was administered to 120 neurologically intact and 89 neurologically affected subjects, yielding data from 43350 individual movements.

This objective methodology to identify a core set of validated metrics based on pathophysiological hypotheses and quantitative selection criteria can complement currently applied paradigms for selecting digital health metrics [17, 24– 28, 38]. While consensus-based recommendations from groups of experts are indispensable for constructing high-level hypothesis (e.g., which body functions to assess in a given context), the selection of specific sensor-based metrics should solely be implemented based on objective and data-driven evaluation criteria to avoid selection bias. Also, guidelines to pool data within systematic reviews, often intended for the selection of conventional assessments, need to be considered carefully in the context of digital health metrics. Compared to conventional assessments that often provide a single, intuitively understandable, task-specific metric (e.g., FMA-UE score), a plethora of abstract digital health metrics exists and the same metric (e.g., *log jerk*) can be extracted from all technologies sharing similar sensor data. However, for a meaningful interpretation of sensor-based metrics, it is essential to consider them in light of the assessment context, as data processing steps (e.g., filter design), assessment platform type (e.g., end-effector or camera-based system), task type (e.g,. goal-directed or explorative movements), and target population (e.g, neurological or muscoskeletal impairments) strongly influences the anticipated hypotheses and clinimetric properties [13]. This emphasizes the importance of a validation of each metric in its specific context (i.e., assessment platform, task, and target population). While objective metric selection algorithms leveraging on the nowadays existing big data sets are already well established in the machine learning domain (therein referred to as *feature selection algorithms*) [24], these usually rely on accurate ground truth (i.e., supervised learning) about the targeted impairment, which is unfortunately often not available in the healthcare domain. Hence, the proposed approach should be seen as an unsupervised metric selection framework aimed to provide a solid foundation of evidence that is required to better transfer research findings into clinical healthcare environments [20, 35].

### 4.1 Specific methodological contributions

In line with literature [39, 40, 57], the mixed effect model analysis (Table SM2) revealed that a high amount of all metrics were significantly modulated by age (69.7% of all metrics) and sex (44.7%), whereas a more selective influence was found for tested side (27.6%) and hand dominance (6.6%). For an accurate assessment of sensorimotor disability without confounds, it is therefore essential to account for these factors when comparing between neurologically intact and affected subjects with different demographics. The presented analysis adds an important methodological contribution to previous work that used linear models to compensate for confounds by additionally evaluating the quality of these models [37, 107–109]. This allowed to discard metrics for which the confounds could not be accurately modeled (16.8% of all metrics). Especially metrics that have mathematical support with two finite boundaries (e.g., 0% and 100%) received low model quality, which can result from skewness and heteroscedasticity that can not be corrected using variance-stabilizing transformations, such as the Box-Cox method. Such metrics should therefore be considered carefully and other modeling approaches, for example based on beta distributions, might be required to accurately compensate for the effect of measurement confounds [110].

Eighty-three percent of all metrics (Table 1) were discarded through the second selection step. It is fundamental to understand that these evaluation criteria (AUC, ICC, SRD%, *η*) are complementary to each other, focusing on different components of intra-subject and inter-subject variability, which are all essential to sensitively monitor impairments. It is therefore not sufficient to solely consider a subset of these criteria, as often done in literature. Evaluating the validity of sensor-based metrics using a *reference population* and ROC analysis is superior to the more commonly applied correlations with conventional scales (concurrent validity) [21, 22]. A reason for this is that sensor-based approaches are being expected to provide complementary information to conventional scales that improves upon their limitations, thereby challenging the definition of accurate hypothesis about the correlation between conventional and sensor-based scales. Nevertheless, comparisons between metrics and conventional scales can help to better interpret sensor-based metrics or to test their sensitivity to impairment severity, as attempted in the last validation step. This analysis was not used as a criteria for metric selection as, to expect trends across subgroups, each sensor-based metric would require a carefully selected clinical counterpart that captures a similar physiological construct. Also, stepwise regression approaches that model conventional scales in order to select metrics have been extensively applied even though they have been considered bad practice due to statistical shortcomings [111–114].

Lastly, a simulated metric without relevant information content (*simulated Gaussian noise*) was rejected in the first and second selection step, thereby providing evidence that the framework allows to discard certain physiologically-irrelevant metrics.

### 4.2 A core set of validated metrics for the VPIT

Applying the proposed framework, ten almost independent metrics (Table 3) were identified as a validated core set for the VPIT and were able to reliably assess the severity of multiple sensorimotor impairments in arm and hand for subjects with mild to moderate disability levels (i.e., the target population of the VPIT). These metrics were related to the movement characteristics smoothness, efficiency, speed, endpoint error, and grip force coordination during specific phases of the task (*transport*, *return*, *peg approach*, *hole approach*). While these characteristics are generally expected to inform on abnormal feedforward control, impaired somatosensory feedback, increased muscle tone, abnormal flexor synergies, dysmetria, and weakness, the clustering of the metrics into five factors allows to further speculate about their interpretation (Table 3). The first factor was dominated by movement efficiency metrics (*path length ratio transport* and *return*), and the *jerk peg approach* as a descriptor for the endpoint error of a movement, thereby fully characterizing the speed-accuracy tradeoff that is a typical characteristic of goal-directed movements [51, 63]. The second factor contained metrics focusing on movement quality (smoothness) during *transport* and *return*, which is expected to describe impaired feedforward control of arm movements. Hence, it is unlikely that the first factor also informs on feedforward control. We therefore expect the movement efficiency metrics (first factor) to be rather related to flexor synergy patterns, weakness, proprioceptive deficits, and dysmetria. Among these impairments, weakness and proprioceptive deficits are most commonly observed in neurological disorders [2, 115]. The third factor focused on grip force coordination during *transport* (*grip force rate num. peaks transport* and *grip force rate spectral arc length transport*), which is expected to be related to abnormal feedforward control and impaired somatosensory feedback. The dissociation between factor one and three is interesting, as it suggests different control schemes underlying the regulation of arm movements and grip forces. A tight predictive coupling between the modulation of grip forces and rapid arm movements has been reported in neurologically intact subjects [116]. The factor analysis suggests that this predictive coupling might possibly be disrupted in neurologically affected subjects, potentially due to altered sensory feedback (e.g., proprioception) leading to inaccurate predictive internal models or abnormal neural transmission (e.g., corticospinal tract integrity) [47, 50]. Reduced corticospinal tract integrity can also lead to weakness and could affect movement speed, as described by factor four (*velocity max. return*) [47]. This factor might further be influenced by an altered inhibition of the supraspinal pathways, often resulting from upper motor neuron lesions, leading to increased muscle tone and thereby altered movement speed [65]. Lastly, the fifth factor covered grip force coordination during *hole approach*, thereby diverging from the coordination of grip forces during gross movements (*transport*) as described by factor 3 and focusing more on grip force coordination during precise position adjustments. This suggests that the two phases are differently controlled, potentially because the *hole approach* is more dominated by sensory and cognitive feedback loops guiding the precise insertion of the peg, whereas gross movements (*transport*) are more dominated through feedforward mechanisms [50]. Also, the physiological control origin of the two movement phases might differ, as gross movements are expected to be orchestrated by the reticulospinal tract, whereas precise control are more linked to the corticospinal tract [117].

Even though the *task completion time* did not pass the selection procedure due to strong learning effects, one might still consider to report this metric when using the VPIT in a cross-sectional manner as its intuitive interpretation allows to give an insightful first indication about the overall level of impairment that might potentially be interesting for both clinical personnel and the tested patient.

The added clinical value of the VPIT core metrics compared to existing conventional assessments is visible in Figure 4 and 5, as the former allowed to detect sensorimotor impairments in certain subjects that did not show any deficits according to the typically used conventional scales. Such a sensitive identification of sensorimotor impairments might allow to provide evidence for the potential of additional neurorehabilitation. Further, the identified core set of metrics can efficiently inform on multiple impairments, both sensory and motor, in arm and hand with a single task that can typically be performed within 15 minutes. This advances the state-of-the-art that mainly focused on the evaluation of arm movements [14, 57, 118], or required more complex or time-consuming measurement setups (e.g., optical motion capture) to quantify arm and hand movements while also neglecting grasping function [119]. Such a fine-grained evaluation covering multiple sensorimotor impairments can help to stratify subjects into homogeneous groups with low inter-subject variability. This is important to reduce the required number of subjects to demonstrate significant effects of novel therapies in clinical trials [14].

### 4.3 Limitations and future directions

The proposed methodology should be considered in light of certain limitations. Most importantly, the framework was especially designed for metrics aimed at longitudinally monitoring impairments and might need additional refinement when transferring it to other healthcare applications (e.g., screening of electronic health record data) with different clinical requirements or data types. Additionally, the definition of multiple cut-off values for the metric selection process influences the final core set of metrics. Even though most of the cut-offs were based on accepted definitions from the research community (e.g., COSMIN guidelines), we acknowledge that the optimality of these values needs to be further validated from a clinical point of view. To evaluate measurement error and learning effects, novel cut-offs were introduced based on the distribution of observed values for the VPIT with the goal to exclude metrics that showed highest measurement error and strongest learning effects. It is important to note that this only considers the relative and not the absolute level of measurement error. However, this can only be adequately judged using data recorded pre- and post-intervention, allowing to compare the measurement error (SRD%) to intervention-induced physiological changes (minimal important clinical difference) [22]. Hence, the rather high absolute level of observed measurement errors for the VPIT (up to 57.7% of the range of observed values) warrants further critical evaluation with longitudinal data. Also, it is important to note that, even though certain VPIT-based metrics did not pass the selection procedure, they might still prove to be valid and reliable for other assessment tasks and platforms, or more specific subject populations. In this context, it should be stressed that test-retest reliability, measurement error, and learning effects for the VPIT were evaluated with neurologically intact subjects and might require additional investigation in neurological populations.

## 5 Conclusions

We proposed a data-driven framework for selecting and validating digital health metrics based on the targeted impairments, the influence of participant demographics, and their clinimetric properties. In a use-case with the VPIT, the methodology enabled the selection and validation of a core set of ten kinematic and kinetic metrics out of 77 initially proposed metrics. The chosen metrics were able to accurately describe the severity of multiple sensorimotor impairments in a cross-sectional manner and have high potential to sensitively monitor neurorehabilitation and to individualize interventions. Additionally, an in-depth physiological motivation of these metrics and the interpretation based on an exploratory factor analysis allowed to better understand their relation to the targeted impairments. Hence, this work makes an important contribution to implement digital health metrics as complementary endpoints for clinical trials and routine, next to the still more established conventional scales and patient reported metrics [120]. We urge researchers and clinicians to capitalize on the promising properties of digital health metrics and further contribute to their validation and acceptance, which in the long-term will lead to a more thorough understanding of disease mechanisms and enable novel applications (e.g., personalized predictions of rehabilitation outcomes) with the potential to improve healthcare quality.

## Ethics approval and consent to participate

All experimental procedure were approved by the responsible local Ethic Committees (approval numbers in SM) and all subjects gave written informed consent prior to participation in the experiments.

## Consent for publication

Not applicable.

## Competing interests

The authors declare no competing interests.

## Availability of data and materials

The datasets used and/or analysed during the current study are available from the corresponding author upon reasonable request.

## Funding

This project received funding from the European Union’s Horizon 2020 research and innovation programme under grant agreement No. 688857 (SoftPro), from the Swiss State Secretariat for Education, Research and Innovation (15.0283-1), from the P&K Puhringer Foundation, by the James S. McDonnell Foundation (90043345, 220020220), and by the Canadian Institutes of Health Research in partnership with the Fondation de l’Ataxie Charlevoix-Saguenay (Emerging Team Grant TR2-119189). CG holds a career-grant-funding from Fonds de recherche en santé du Québec (22193, 31011).

## Contributions

Study design: CK, AS, IL, CG, JH, PF, ARL, RG, OL. Data collection: CMK, AS, JH, CG, IL, OL. Data analysis: CMK. Data interpretation: CMK, MDR, RG, OL. Manuscript writing: CMK, RG, OL. Manuscript review: MDR, AS, IL, JH, PF, RG, OL. All authors read and approved the final manuscript.

## Acknowledgements

The authors thank Marie-Christine Fluet, Sascha Motazedi Tabrizi, Werner Popp, Joachim Cerny, Isabelle Lessard, Caroline Lavoie, and Meret Branscheidt for help during data collection and the insightful discussions.

## 1 Supplementary Material: Methods

### 1.1 Participants

Neurologically intact subjects were recruited at ETH Zurich (Zurich, Switzerland). Stroke patients were tested at the University Hospital of Zurich (Zurich, Switzerland), the cereneo Center for Neurology and Rehabilitation (Vitznau, Switzerland), and the Zentrum für ambulante Rehabilitation (ZAR, Zurich, Switzerland) as part of the Study of Motor Learning and Acute Recovery Time Course in Stroke (SMARTS) or the synergy-based open-source foundations and technologies for prosthetics and rehabilitation (SoftPro). Multiple sclerosis (MS) patients were recruited at Hasselt University (Hasselt, Belgium) and at the Rehabilitation and MS Center Overpelt (Overpelt, Belgium), some of them as part of the individualised technology-supported and robot-assisted virtual learning environment(ITRAVLE) study. Exclusion criteria involved the inability to lift the arm against gravity, to flex/extend the fingers, and the presence of any concomitant disease affecting the upper limb. The studies involving stroke patients additionally used increased muscle tone, severe sensory deficits, hemorrhagic infarct, traumatic brain injury as exclusion criteria. MS patients had to be diagnosed according to the McDonald criteria. All clinical assessments were performed within the same or few days of the Virtual Peg Insertion Test (VPIT) assessment. Experimental procedures were approved by the local Ethics Committees: neurologically intact subjects subjects: EK2010-N-40; stroke patients: EKNZ-2016-02075, KEK-ZH 2011-0268; MS patients: CME2013/314, ML9521 (S55614), B322201318078; ARSACS patients: 2012-012.

## 2 Supplementary Material: Results

The metrics that did not fullfil the required quality of the models, according to the C1 and C2 criteria, were *spectral arc length transport*, *number of velocity peaks transport*, *distance to max. velocity transport*, *time to max. velocity transport*, *number of velocity peaks return*, *throughput transport*, *initial movement angle transport θ*_1_, *initial movement angle θ*_2_, *collision force max. return*, *grip force rate number of peaks buildup*, *grip force rate spectral arc length buildup*, *grip force rate number of peaks release*, and *simulated Gaussian noise*.

---

**Table SM1:** Detailed demographics and clinical information for each body side of each included neurologically impaired subject.

| Disease | Age<br>(yrs) | Sex | Tested side | Affected side | Dominant side | Chronicity<br>(yrs) | FMA-UE<br>(0-66) | ARAT<br>(0-57) | NHPT<br>(s) | EDSS<br>(0-10) |
| --- | --- | --- | --- | --- | --- | --- | --- | --- | --- | --- |
| Stroke | 67 | Male | Right | Left | Right | 2.09 | 66 | 57 | 23.25 | - |
| Stroke | 55 | Male | Left | Left | Right | 1.69 | 54 | 56 | 33.25 | - |
| Stroke | 55 | Male | Right | Left | Right | 1.69 | 66 | 57 | 21.85 | - |
| Stroke | 55 | Male | Left | Right | Right | 2.01 | 65 | 57 | 22.82 | - |
| Stroke | 55 | Male | Right | Right | Right | 2.01 | 49 | 55 | 29.28 | - |
| Stroke | 52 | Male | Left | Left | Right | 2.74 | 55 | 52 | 35.36 | - |
| Stroke | 52 | Male | Right | Left | Right | 2.74 | 65 | 57 | 20.99 | - |
| Stroke | 73 | Male | Left | Right | Right | 0.89 | 62 | - | - | - |
| Stroke | 69 | Female | Right | Left | Right | 0.86 | 61 | 57 | 20.32 | - |
| Stroke | 67 | Male | Left | Left | Right | 2.42 | 50 | - | - | - |
| Stroke | 67 | Male | Right | Left | Right | 2.42 | 66 | - | - | - |
| Stroke | 40 | Female | Left | Right | Right | 0.77 | 56 | 45 | - | - |
| Stroke | 40 | Female | Right | Right | Right | 0.77 | 49 | 49 | - | - |
| Stroke | 71 | Male | Left | Left | Left | 4.49 | 40 | 35 | 196.69 | - |
| Stroke | 71 | Male | Right | Left | Left | 4.49 | 65 | 57 | 15.03 | - |
| Stroke | 59 | Female | Left | Left | Right | 4.35 | 50 | 47 | 17.70 | - |
| Stroke | 59 | Female | Right | Left | Right | 4.35 | 66 | 57 | 12.57 | - |
| Stroke | 88 | Female | Left | Left | Right | 1.65 | 37 | 39 | 42.17 | - |
| Stroke | 88 | Female | Right | Left | Right | 1.65 | 63 | - | 14.33 | - |
| Stroke | 69 | Female | Left | Right | Right | 0.58 | 63 | 57 | 19.81 | - |
| Stroke | 69 | Female | Right | Right | Right | 0.58 | 44 | 39 | 49.16 | - |
| Stroke | 59 | Female | Left | Right | Right | 1.94 | 66 | 57 | 21.50 | - |
| Stroke | 59 | Female | Right | Right | Right | 1.94 | 57 | 56 | 21.63 | - |
| Stroke | 50 | Female | Right | Left | Right | 4.83 | 64 | - | - | - |
| Stroke | 61 | Male | Left | Right | Right | 8.70 | 66 | 56 | 24.51 | - |
| Stroke | 61 | Male | Right | Right | Right | 8.70 | 38 | 42 | 34.95 | - |
| Stroke | 59 | Male | Left | Left | Right | 1.64 | 46 | 40 | 40.84 | - |
| Stroke | 59 | Male | Right | Left | Right | 1.64 | 63 | 57 | 14.85 | - |
| Stroke | 69 | Male | Left | Left | Right | 0.51 | 53 | 51 | 23.08 | - |
| Stroke | 69 | Male | Right | Left | Right | 0.51 | 63 | 56 | 13.67 | - |
| Stroke | 55 | Male | Left | Left | Right | 1.45 | 59 | 57 | 28.08 | - |
| Stroke | 55 | Male | Right | Left | Right | 1.45 | 66 | 57 | 18.50 | - |
| Stroke | 42 | Male | Left | Left | Right | 0.48 | 39 | 30 | - | - |
| Stroke | 42 | Male | Right | Left | Right | 0.48 | 65 | 57 | 20.47 | - |
| Stroke | 51 | Female | Left | Right | Right | 0.97 | 66 | 57 | 21.01 | - |
| Stroke | 51 | Female | Right | Right | Right | 0.97 | 61 | 57 | 25.70 | - |
| Stroke | 58 | Male | Left | Right | Right | 0.48 | 62 | 57 | 23.33 | - |
| Stroke | 58 | Male | Right | Right | Right | 0.48 | 42 | 53 | 26.00 | - |
| Stroke | 46 | Male | Left | Left | Right | 1.05 | 57 | 42 | 24.03 | - |
| Stroke | 46 | Male | Right | Left | Right | 1.05 | 66 | 57 | 23.09 | - |
| Stroke | 76 | Male | Left | Right | Right | 2.74 | 66 | 55 | 39.73 | - |
| Stroke | 76 | Male | Right | Right | Right | 2.74 | 60 | 54 | 29.19 | - |
| Stroke | 53 | Female | Left | Right | Right | 2.98 | 66 | 57 | 22.99 | - |
| Stroke | 53 | Female | Right | Right | Right | 2.98 | 58 | 55 | 20.67 | - |
| Stroke | 62 | Male | Left | Right | Right | 14.65 | 66 | 57 | 19.58 | - |
| Stroke | 62 | Male | Right | Right | Right | 14.65 | 34 | 33 | 154.00 | - |
| Stroke | 62 | Male | Left | Right | - | - | - | 57 | 24.60 | - |
| Stroke | 62 | Male | Right | Right | - | - | - | 43 | 86.00 | - |
| Stroke | 54 | Female | Left | Left | Right | 1.00 | 66 | 57 | - | - |
| Stroke | 54 | Female | Right | Left | Right | 1.00 | 66 | 57 | - | - |
| Stroke | 67 | Male | Left | Left | Right | 0.46 | 66 | 57 | - | - |
| Stroke | 67 | Male | Right | Left | Right | 0.46 | 66 | 57 | - | - |
| Stroke | 52 | Male | Left | Left | Right | 0.23 | 66 | 57 | - | - |
| Stroke | 52 | Male | Right | Left | Right | 0.23 | 66 | 57 | - | - |
| Stroke | 46 | Male | Left | Right | Right | 0.23 | 66 | 57 | - | - |
| Stroke | 71 | Male | Left | Left | Right | 0.23 | 64 | 57 | - | - |
| Stroke | 71 | Male | Right | Left | Right | 0.23 | 66 | 57 | - | - |
| Stroke | 48 | Male | Left | Right | Right | 0.02 | 57 | 57 | - | - |
| Stroke | 48 | Male | Right | Right | Right | 0.02 | 66 | 47 | - | - |
| Stroke | 45 | Female | Right | Left | Right | 0.02 | 66 | 57 | - | - |
| Stroke | 55 | Female | Right | Left | Right | 0.08 | 66 | 57 | - | - |
| Stroke | 65 | Male | Left | Left | Right | 0.23 | 60 | - | - | - |
| Stroke | 65 | Male | Right | Left | Right | 0.02 | 62 | 53 | - | - |
| Stroke | 43 | Male | Left | Right | Right | 0.46 | 66 | 57 | - | - |
| Stroke | 43 | Male | Right | Right | Right | 0.46 | 66 | 56 | - | - |
| Stroke | 41 | Female | Right | Left | Right | 0.02 | 64 | - | - | - |
| Stroke | 35 | Male | Left | Left | Right | 0.46 | 61 | 57 | - | - |
| Stroke | 35 | Male | Right | Left | Right | 0.02 | 64 | 57 | - | - |
| Stroke | 76 | Male | Right | Left | Left | 0.23 | 66 | 57 | - | - |
| Stroke | 86 | Male | Right | Left | Right | 0.02 | 62 | 56 | - | - |
| Stroke | 50 | Male | Left | Left | Right | 1.00 | 65 | 57 | - | - |
| Stroke | 49 | Male | Right | Left | Left | 0.23 | 66 | 57 | - | - |
| Stroke | 74 | Male | Left | Right | Right | 0.02 | 66 | 57 | - | - |
| Stroke | 81 | Female | Left | Right | Right | 0.23 | 66 | 57 | - | - |
| Stroke | 65 | Female | Left | Left | Right | 0.23 | 66 | 56 | - | - |
| Stroke | 65 | Female | Right | Left | Right | 0.23 | 66 | 57 | - | - |
| Stroke | 21 | Male | Left | Right | Right | 0.02 | 63 | 57 | - | - |
| Stroke | 21 | Male | Right | Right | Right | 0.02 | 66 | 56 | - | - |
| Stroke | 87 | Female | Left | Right | Left | 0.02 | 66 | 57 | - | - |
| Stroke | 87 | Female | Right | Right | Left | 0.02 | 50 | 29 | - | - |
| Stroke | 54 | Male | Left | Right | Left | 0.46 | 66 | 57 | - | - |
| Stroke | 54 | Male | Right | Right | Left | 0.46 | 54 | 57 | - | - |
| Stroke | 57 | Male | Left | Left | Right | 0.02 | 66 | 57 | - | - |
| Stroke | 57 | Male | Right | Left | Right | 0.02 | 61 | 57 | - | - |
| Stroke | 70 | Female | Right | Left | Right | 0.53 | 66 | - | 16.46 | - |
| Stroke | 57 | Male | Right | Right | - | 0.48 | 66 | - | 22.61 | - |
| Stroke | 73 | Male | Left | Left | Right | 0.53 | 63 | - | 28.55 | - |
| Stroke | 56 | Male | Left | Left | Right | 0.03 | 25 | - | 60.81 | - |
| Stroke | 63 | Male | Left | Right | Right | 0.48 | 66 | - | 14.33 | - |
| ARSACS | 41 | Female | Left | Both | Right | - | - | - | 28.59 | - |
| ARSACS | 41 | Female | Right | Both | Right | - | - | - | 37.14 | - |
| ARSACS | 29 | Male | Left | Both | Right | - | - | - | 56.98 | - |
| ARSACS | 29 | Male | Right | Both | Right | - | - | - | 40.34 | - |
| ARSACS | 56 | Female | Left | Both | Left | - | - | - | 83.59 | - |
| ARSACS | 56 | Female | Right | Both | Left | - | - | - | 95.20 | - |
| ARSACS | 37 | Male | Left | Both | Left | - | - | - | 36.36 | - |
| ARSACS | 37 | Male | Right | Both | Left | - | - | - | 46.72 | - |
| ARSACS | 26 | Female | Left | Both | Right | - | - | - | - | - |
| ARSACS | 26 | Female | Right | Both | Right | - | - | - | - | - |
| ARSACS | 37 | Female | Left | Both | Right | - | - | - | - | - |
| ARSACS | 37 | Female | Right | Both | Right | - | - | - | - | - |
| ARSACS | 31 | Male | Left | Both | Right | - | - | - | 29.88 | - |
| ARSACS | 31 | Male | Right | Both | Right | - | - | - | 23.52 | - |
| ARSACS | 58 | Male | Left | Both | Right | - | - | - | 60.43 | - |
| ARSACS | 58 | Male | Right | Both | Right | - | - | - | 47.33 | - |
| MS | 52 | Female | Left | Both | Right | 29.00 | - | 37 | 45.25 | 7.0 |
| MS | 52 | Female | Right | Both | Right | 29.00 | - | 47 | 24.75 | 7.0 |
| MS | 69 | Male | Right | Both | Right | 19.00 | - | 44 | 140.27 | 7.5 |
| MS | 25 | Female | Left | Both | Right | 6.00 | - | 52 | 29.35 | 6.0 |
| MS | 25 | Female | Right | Both | Right | 6.00 | - | 53 | 29.62 | 6.0 |
| MS | 42 | Female | Left | Both | Right | 1.00 | - | 56 | 27.81 | 4.0 |
| MS | 42 | Female | Right | Both | Right | 1.00 | - | 54 | 20.48 | 4.0 |
| MS | 59 | Female | Left | Both | Left | 5.00 | - | 49 | 27.76 | 7.0 |
| MS | 56 | Female | Left | Both | Right | 10.00 | - | 49 | 33.72 | 7.0 |
| MS | 56 | Female | Right | Both | Right | 10.00 | - | 29 | 89.79 | 7.0 |
| MS | 65 | Male | Left | Both | Left | 19.00 | - | 52 | 39.90 | 8.0 |
| MS | 63 | Female | Left | Both | Right | 8.00 | - | 57 | 20.84 | 4.5 |
| MS | 63 | Female | Right | Both | Right | 8.00 | - | 54 | 35.04 | 4.5 |
| MS | 76 | Female | Left | Both | Right | 38.00 | - | 43 | 27.01 | 5.0 |
| MS | 76 | Female | Right | Both | Right | 38.00 | - | 34 | 34.46 | 5.0 |
| MS | 60 | Male | Left | Both | Right | 21.00 | - | 52 | 31.48 | 7.0 |
| MS | 60 | Male | Right | Both | Right | 21.00 | - | 53 | 25.29 | 7.0 |
| MS | 42 | Female | Right | Both | Right | 21.00 | - | 39 | 74.39 | 7.5 |
| MS | 46 | Male | Left | Both | Right | 11.00 | - | 55 | 30.58 | 5.5 |
| MS | 46 | Male | Right | Both | Right | 11.00 | - | 56 | 23.23 | 5.5 |
| MS | 70 | Female | Left | Both | Right | 37.00 | - | 53 | 29.86 | 6.0 |
| MS | 70 | Female | Right | Both | Right | 37.00 | - | 45 | 53.21 | 6.0 |
| MS | 36 | Female | Right | Both | Right | 6.76 | 61 | 56 | 22.87 | 7.5 |
| MS | 40 | Male | Right | Both | Left | 12.55 | 53 | 44 | 56.17 | 7.5 |
| MS | 35 | Male | Left | Both | Right | 0.97 | 65 | 57 | 22.90 | 4.5 |
| MS | 35 | Male | Right | Both | Right | 0.97 | 65 | 52 | 24.90 | 4.5 |
| MS | 52 | Female | Left | Both | Right | 9.66 | 62 | 56 | 35.13 | 5.5 |
| MS | 52 | Female | Right | Both | Right | 9.66 | 65 | 56 | 29.31 | 5.5 |
| MS | 65 | Female | Right | Both | Both | 14.48 | 61 | 52 | 55.37 | 7.5 |
| MS | 53 | Male | Right | Both | Right | 9.66 | 62 | 56 | 23.93 | 2.5 |
| MS | 59 | Male | Left | Both | Right | 1.93 | 63 | 56 | 28.70 | 4.0 |
| MS | 59 | Male | Right | Both | Right | 1.93 | 63 | 55 | 47.49 | 4.0 |
| MS | 35 | Female | Left | Both | Right | 14.48 | 62 | 51 | 50.17 | 7.5 |
| MS | 38 | Male | Left | Both | Right | 2.90 | - | - | - | 3.5 |
| MS | 38 | Male | Right | Both | Right | 2.90 | - | - | - | 3.5 |
| MS | 66 | Female | Left | Both | Left | 16.42 | - | - | - | 7.5 |
| MS | 66 | Female | Right | Both | Left | 16.42 | - | - | - | 7.5 |
| MS | 22 | Male | Left | Both | Right | 3.86 | - | - | - | 6.5 |
| MS | 22 | Male | Right | Both | Right | 3.86 | - | - | - | 6.5 |
| MS | 38 | Female | Left | Both | Right | 8.69 | - | - | - | 7.0 |
| MS | 38 | Female | Right | Both | Right | 8.69 | - | - | - | 7.0 |
| MS | 61 | Male | Left | Both | Right | 5.79 | - | - | 20.00 | 5.0 |
| MS | 61 | Male | Right | Both | Right | 5.79 | - | - | 26.27 | 5.0 |
| MS | 63 | Female | Right | Both | Right | 6.76 | - | - | 28.00 | 6.0 |
| MS | 63 | Male | Right | Both | Right | 29.93 | - | - | 16.00 | 3.0 |

**Table SM2:**
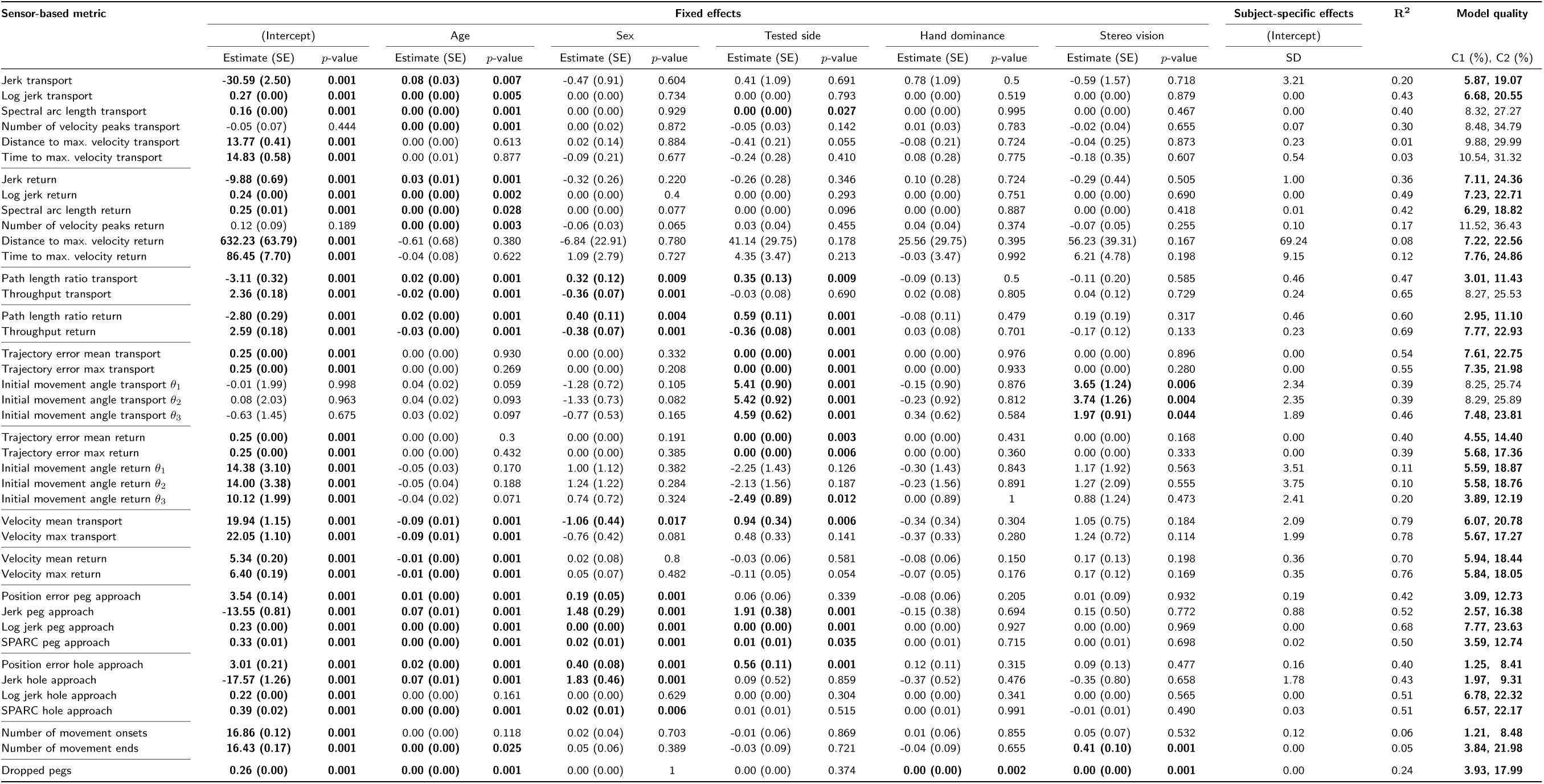
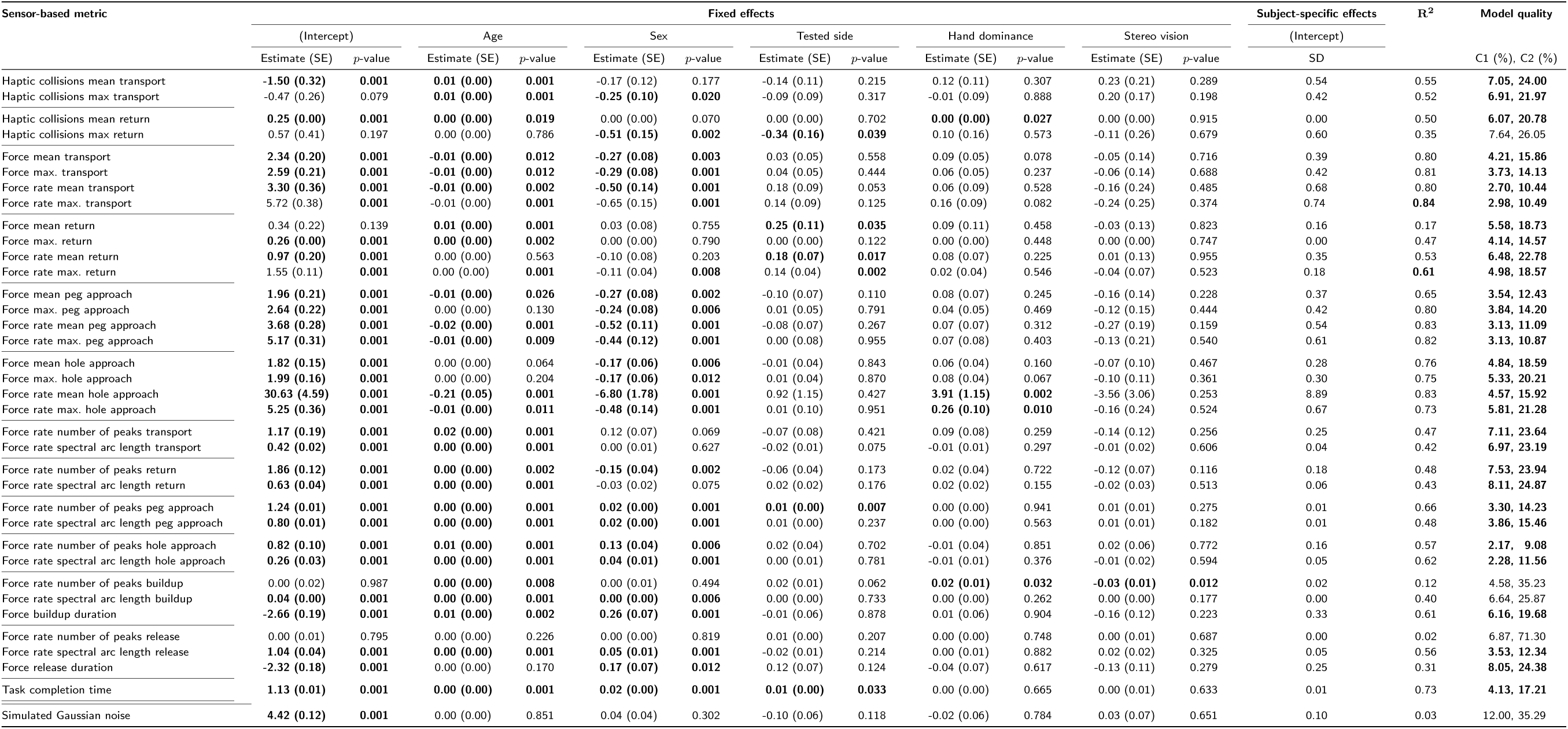
Influence of potential confounds on each sensor-based metric. For each metric, a mixed effect model was fitted to the Box-Cox-transformed outcome measure. Bold entries indicate that the fixed effect contributed in a statistically significant manner to model quality according to a simulated likelihood ratio test or that model quality was at least moderate according to the criteria *C*1 and *C*2. Abbreviations: SE: standard error; SD: standard deviation. *R*^2^: adjusted coefficient of determination. SPARC: spectral arc length.

**Figure SM1:**
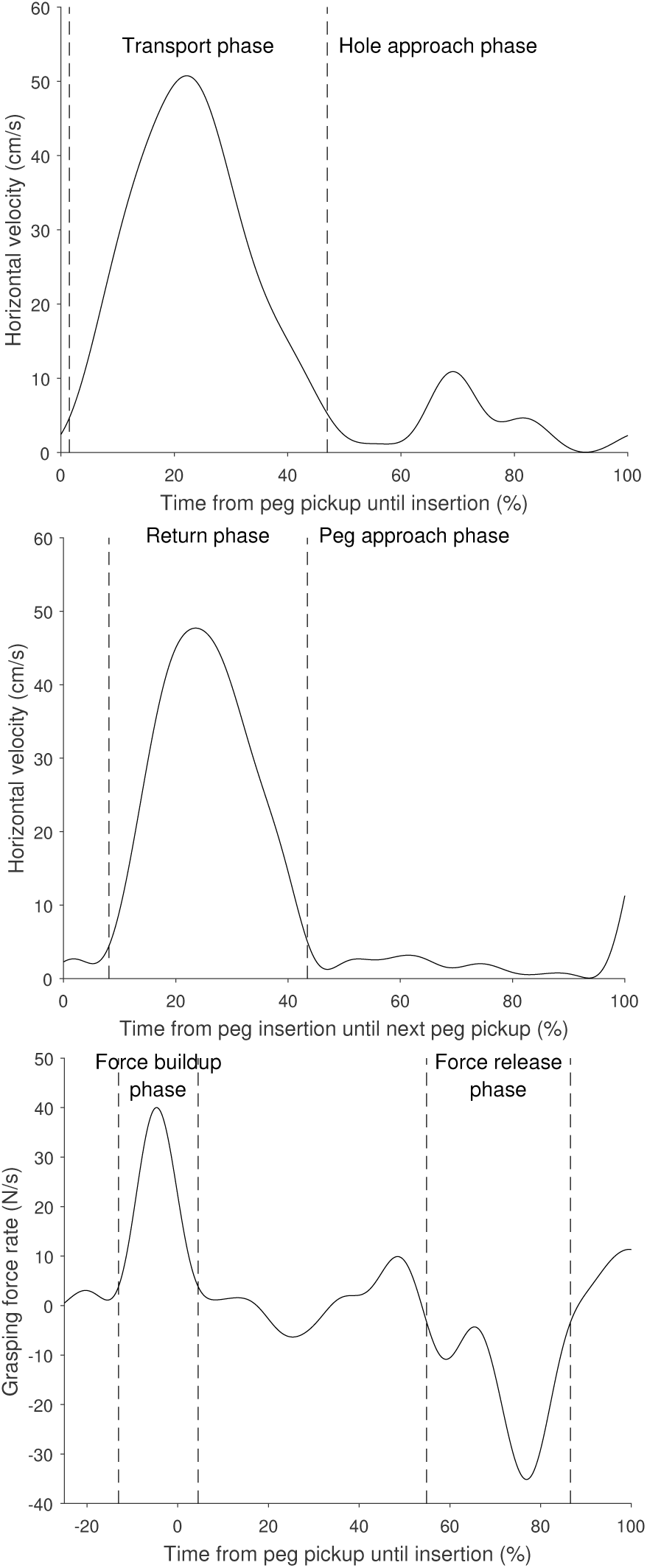
Temporal segmentation of kinematic and kinetic trajectories. Representative example from one neurologically intact subject (49yrs, female, tested hand left, dominant hand right).

**Figure SM2:**
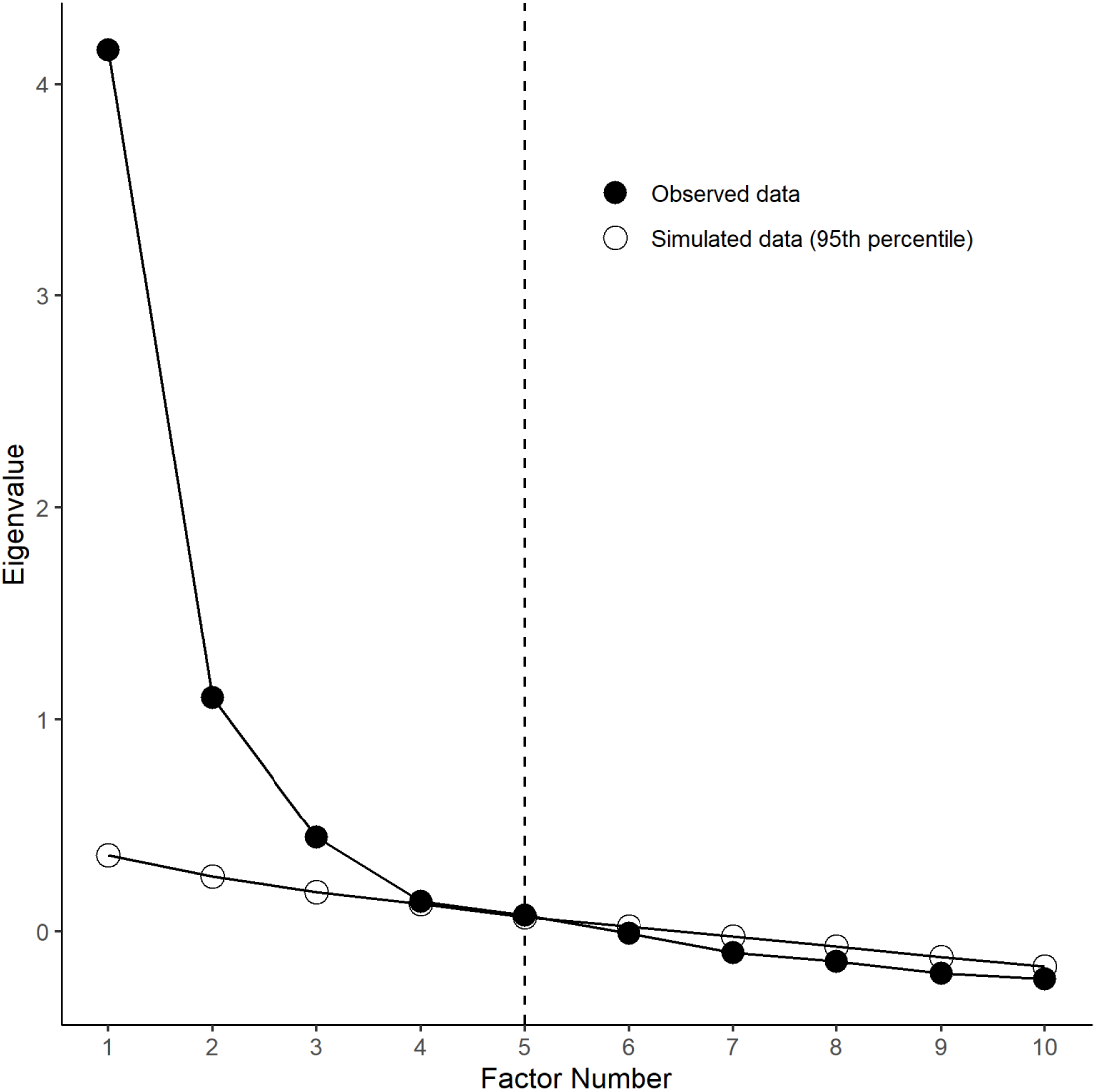
Scree plot for estimating the number of latent variables *k* in the factor analysis. Parallel analysis was used to simulate a lower bound of an eigenvalues magnitude that each eigenvalue in the observed data needs to fulfill. The chosen number of factors was set to five accordingly.

